# Force Volume Atomic Force Microscopy of THP-1 and Primary Mouse Macrophages Reveals that the Mechanical Signature of Macrophage Polarization is Model-Dependent

**DOI:** 10.64898/2026.11.17.767239

**Authors:** Beatriz Cantero, Mauricio Cano, Lara Aiassa, Claudia Codano, Sophie Strawbridge, Loris Rizzello, Giuseppe Battaglia, Gabriel Gomila, Annalisa Calò

**Author notes:** E-mail:* Beatriz Cantero; Mauricio Cano; Lara Aiassa; Claudia Codano; Sophie Strawbridge; Loris Rizzello; Giuseppe Battaglia. Corresponding authors: Gabriel Gomila, Annalisa Calò *E-mail:* Gabriel Gomila; Annalisa Calò.

## Abstract

Macrophages are immune cells with a rich morphological variety and markedly different biological functions when differentiated into M0 (non-activated) macrophages or polarized into M1 (pro-inflammatory) and M2 (anti-inflammatory) phenotypes. Atomic force microscopy (AFM) studies have recently associated a mechanical dimension with macrophage polarization. Herein, the elastic properties of the three phenotypes are measured and compared by force volume AFM in two less explored models: THP-1-derived macrophages (cell line) and mouse bone marrow-derived macrophages (BMDMs, primary cells). High-indentation measurements (indentation ≥ 2 µm) with colloidal probes capture the full cell deformation, and a recently derived spherical model with bottom effect correction provides quantitative values of the Young’s modulus (*E*) that account for substrate effects and cell height with the correct probe geometry. THP-1-derived macrophages show significant mechanical differences, with *E*_M1_ higher than *E*_M2_ and *E*_M0_, whereas BMDMs show no significant differences between phenotypes. Furthermore, BMDM-derived M0 and M2 macrophages show 1.5 to 2 times higher *E* than the corresponding THP-1 phenotypes. The elasticity of cells with different shapes is also evaluated to account for the heterogeneous macrophage morphology found in all samples. These results indicate that the mechanical signature of macrophage polarization is model-dependent.

## 1. Introduction

Macrophages are immune cells characterized by a varied morphology and cellular function both in *in vitro* and *in vivo* models^1,2^. While macrophages are characterized by shades of activation states *in vivo*^3,4^, in cultured samples resting or naïve macrophages (M0 phenotype) can be polarized into phenotypes that represent a rather extreme biological response towards external aggression, i.e. pro-inflammatory (M1) and anti-inflammatory (M2) phenotypes. These two macrophage types have the function of creating inflammation in response to pathogens and restoring the normal tissue homeostasis after inflammation, respectively. The three macrophage phenotypes show marked morphological differences. For example, cultured M0 cells obtained from small non-adherent monocytes^5,6^ typically exhibit a rounded shape or small elongations, while M1 and M2 phenotypes show more extended shapes with pronounced elongations, roughly rounded or with star-like contours^5–7^. These shapes also vary depending on the cell model^8,9^.

Functional characteristics of distinct macrophage phenotypes are assessed through a combination of assays rather than with single specific tests; these include the expression of phenotype-specific surface biomarkers, genetic analysis, cytokine secretion and functional analysis of enzyme activity (inducible nitric oxide synthase (iNOS) and arginase activity)^10–14^. Cell morphological changes upon exposure to polarizing stimuli have also been considered as phenotypic indicators of macrophage polarization^15^. The identification of phenotypic differences in macrophages based on physical rather than biochemical characteristics has been far less explored. Physical cues seem to play an important role in determining macrophage function. For example, in mouse bone marrow-derived macrophages (BMDM) it has been shown that purely physical constraints applied to M0 cells can induce morphological changes, i.e. cell elongation, and consequent polarization to M2^16^. Also, in the same model it was demonstrated that spatial confinement downregulates the proinflammatory characteristics of the M1 phenotype by limiting the area needed for efficient cell activation^17^. Cell mechanics has also been proposed as a separate physical contributor to macrophage function^18,19^. In this context, besides reports employing other single cell approaches^20^, atomic force microscopy (AFM) has emerged as an ideal technique for the mechanical phenotyping of M0, M1 and M2 macrophages. Topographic and mechanical characteristics of living and fixed macrophages from different cell lineages have been studied using nanometric AFM probes^21–24^. More comprehensive elasticity analysis has focused on the live mouse-derived RAW 264.7 macrophage cell line using colloidal probes^25–29^. Live macrophages from patients’ monocytes have also been studied by colloidal probe-based AFM, with the aim of training machine-learning algorithms with physical information (cell elasticity, adhesion, topography and their spatial distribution) for precise, label-free phenotyping^30^. Macrophage phenotypes from other cell models, such as those derived from the monocytic THP-1 cell line and mouse BMDM, which are among the most widely used *in vitro* models to study the biology of the immune system, have been less frequently studied by AFM in live conditions using colloidal probes^25,31^.

For immune cells, cell morphology (volume, surface area) and shape are considered phenomenological determinants of cellular function, often in association with some mechanical property. For example, rounded undifferentiated monocytes show poor adhesion on surfaces for cell culturing, a fact that has been correlated to their circulatory function *in vivo*^32,33^. Vice versa, M1 macrophages, which have a larger surface area, adhere firmly to culture plates, a fact that has been associated with the strong interaction that this highly polarized phenotype has with tissues *in vivo*^34^. Cytoskeletal analysis of macrophages with different shapes also showed differences in the distribution and intensity of main cytoskeleton markers (F-actin, vinculin, α-tubulin)^16,25,30^. Taken together, these observations may indicate macrophage morphology as a possible determinant of macrophage mechanical properties. To our knowledge, there are no reports correlating the heterogeneous macrophage morphology, present even within the same dish after differentiation and polarization protocols, with the corresponding cell mechanics by AFM.

In this work we characterize the morphology and elasticity of THP-1 and mouse BMDM by AFM in high indentation measurements using colloidal probes. For data analysis, we employed a model that we developed for the spherical probe geometry used in the experiments, which includes correction for the cell height and bottom substrate effect^35^. In previous studies using colloidal probes, models accounting for the sample height have been used that consider a paraboloid probe geometry^25–28,30^; in other studies, models that do not include the bottom effect correction have been employed for data analysis^21,23,24^. Both models can introduce inaccuracies in the Young’s modulus determination, when testing thin regions of the cells or when high indentations are applied^35^. To achieve a comprehensive characterization of both cell models, we also analyzed the main cellular shapes present in the samples for each phenotype. Our results show clearly different mechanical behavior of THP-1 and BMDM-derived macrophages, both in terms of the average Young’s modulus and of the phenotypic differences between M0, M1 and M2 cell types.

## 2. Results

In this work, we analyzed a total of 179 cells by force volume AFM, obtained from the THP-1 human monocytic cell line (N = 96 cells) and from the bone marrow of 6-week-old C57BL/6 mice, i.e. BMDM (N = 83 cells). Macrophages of the three different phenotypes were prepared using standard protocols for differentiation that were validated using ELISA, RT-qPCR and flow cytometry (see Supplementary Figures S1–S2 and the Materials and Methods section)^36,37^. Macrophages were cultured on glass-bottom Petri dishes in all cases.

Figure 1 shows optical microscopy images of cells in M0, M1 and M2 samples in the two models. In the case of the THP-1-derived macrophages (Figure 1a), differentiation of monocytes into the M0 phenotype typically produced cells with a rounded shape and, less frequently, cells with small elongations (spindle-like cells). Polarization to M1 and M2 produced cells with progressively more elongated shapes, which were absent in the unpolarized type. Although accurate phenotyping can only be derived from correlative techniques and immunofluorescence^38^, we attribute these morphological characteristics to the development of a different biological function upon polarization and formation of activated states. In M2 samples a more heterogeneous morphology was observed, i.e. branched or dendritic cells were also occasionally present in the THP-1 model, consistent with other findings^6^.

**Figure 1.**
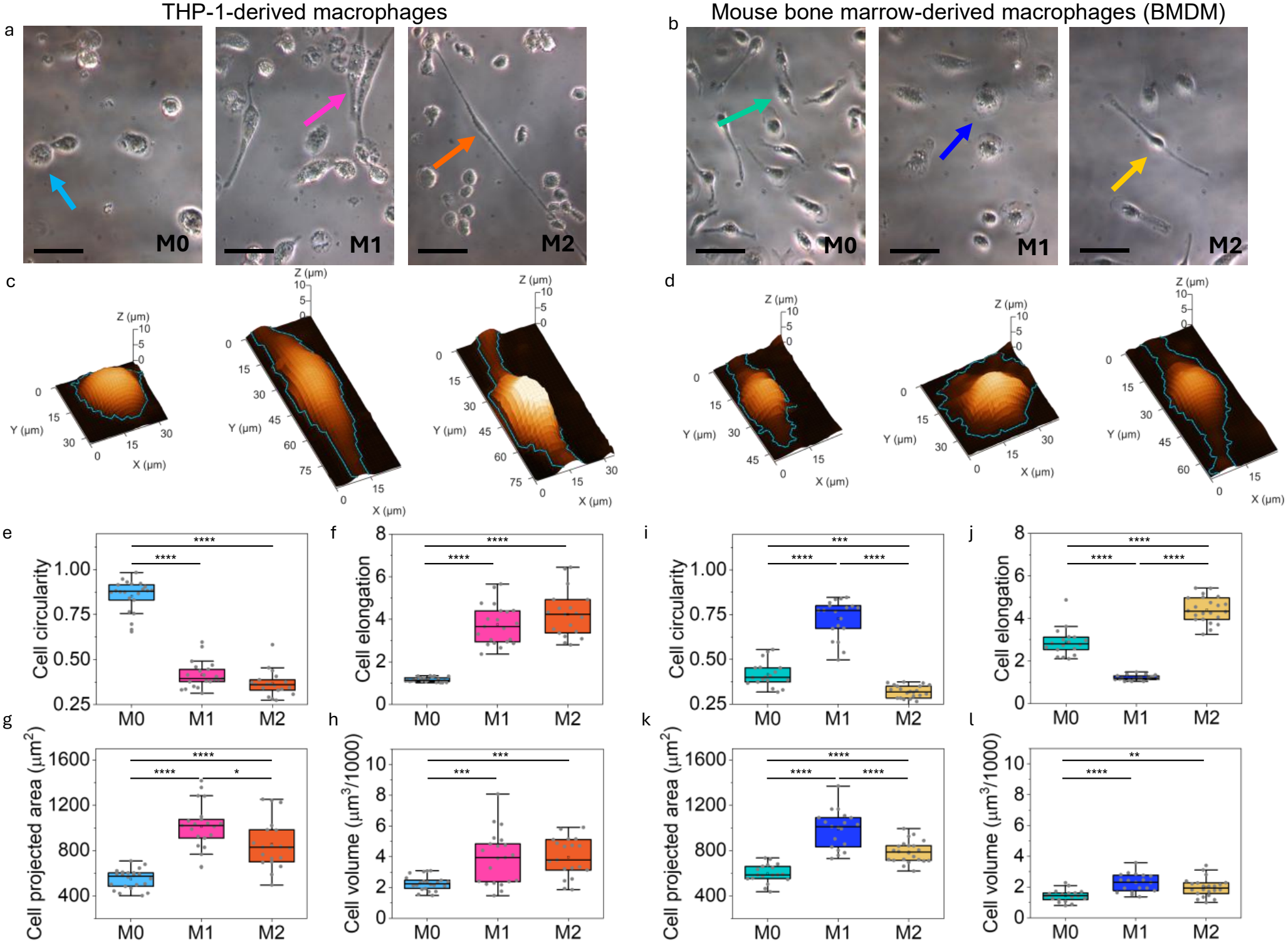
Optical microscopy images of (a) THP-1-derived macrophages and (b) BMDM-derived macrophages, where typical cell shapes associated with the main phenotypes are highlighted with arrows. The scale bar, identical for all optical images, is 50 µm. (c-d) AFM true topographic images (3D view) of representative cells from (a) and (b), respectively, showing the segmentation contour used for the morphometric analysis (blue line). (e–h) Morphometric analysis of THP-1-derived macrophages (N_M0_=22, N_M1_=21, N_M2_=19): cell circularity (e), cell elongation (f), cell projected area (g) and cell volume (h) (see Materials and Methods). (i–l) Same parameters for BMDM (N_M0_=18, N_M1_=19, N_M2_=22). Results of statistical comparison are also shown for each parameter (one-way ANOVA).

Similarly, in the case of the BMDM model, macrophage shape was used as a phenotypic indicator for AFM analysis. After differentiation from bone marrow cells (see Materials and Methods), M0 macrophages showed predominantly a spindle-like shape and exhibited small elongations; upon polarization to M1, cells with a rounded nucleus and extended, roughly circular perinuclear areas were frequently observed, while highly elongated structures were typically present in M2 samples (Figure 1b). Nevertheless, it should be noted that, consistent with other reports using primary macrophages^7,15^, in the BMDM case a richer cell morphology was present in each sample compared to THP-1. For example, cells exhibiting enlarged, roughly rounded cytoplasmic areas were present in all samples, cells with pronounced elongations were present not only in M2 samples but also in M0 samples and cells with small elongations were observed not only in M0 samples, but also in M1 samples. The shapes highlighted with arrows in Figure 1a and 1b are assigned to the distinct macrophage phenotypes in agreement with reported bright field and fluorescence images in the literature^6,7,16^. In the present work, AFM mechanical analysis of all the other cellular shapes encountered in each sample is separately presented for the two models.

Following other reports^25,26,30^, our AFM analysis initially focused on rounded, elongated and highly elongated cells present in M0, M1 and M2 samples respectively (THP-1 model) and cells with small elongations, extended and roughly rounded perinuclear areas, and highly elongated cells present in M0, M1 and M2 samples respectively (BMDM model). Morphometric parameters were obtained from an automatic segmentation of the whole cell area (Otsu thresholding followed by marker-based watershed; see Materials and Methods) using AFM images with accurate height information (true topography, see Figure 1c and 1d), to obtain 2D and 3D cell characteristics. The extracted parameters (cell circularity, cell elongation, cell projected area and cell volume, see Figure 1e-h for the THP-1 model and Figure 1i-l for the BMDM model) well describe the morphological characteristics highlighted in Figure 1a and 1b. The number of cells analyzed of each type is N_M0_=22, N_M1_=21, N_M2_=19 for the THP-1 model and N_M0_=18, N_M1_=19, N_M2_=22 for the BMDM model. Upon polarization of M0 to M1 and M2 in the THP-1 case, a marked decrease in cell circularity and increase in the cell elongation was observed, accompanied by an increase in the cell projected area (Figure 1e-g). The cell volume, which is calculated from the height of each pixel corrected for the substrate level (see Materials and Methods), was higher in the elongated cells of the two polarized phenotypes compared to rounded M0 cells (Figure 1h). In the BMDM case, typical cells of the M1 phenotype clearly exhibited higher circularity, lower elongation and higher projected area compared to typical M0 and M2 cells, confirming the optical observations (Figure 1i-k). The cell volume of the analyzed M1 and M2 cells was higher compared to the M0 cells in the BMDM model (Figure 1l). It is worth noting that in the BMDM model the cell volume was markedly smaller in comparison with the THP-1 model due to a decreased topographic height (see Supplementary Figure S3a and b). Details of the methodology and the equations used to extract the morphological parameters are given in the Materials and Methods section.

Force volume AFM experiments were conducted on each cell. In Figure 2, the workflow used to process the force curves and obtain accurate mechanical and topographic information is shown, using data acquired on an M1 macrophage derived from the THP-1 cell line as an example. Figure 2a shows the ensemble of force curves acquired (17 × 54 force curves). Figure 2b shows the rough topography image, corresponding to the experimental force set point, F_set_=9 nN. Cell and substrate curves were separately analyzed by manual image segmentation (for details on the segmentation used for the morphometric analysis, Figure 1, and AFM mechanical analysis, Figure 2, see the Materials and Methods section). The curves on the hard substrate (N_subs_=46) allowed the determination of the average optical lever sensitivity (OLS) in each map (see Figure 2c that shows the histogram of contact slopes determined, the fitted straight lines and the substrate area used for OLS calibration overlaid on the topography image). This value, together with the cantilever spring constant, separately calibrated using the thermal noise method, was used to obtain force versus z-piezo and force versus tip-sample distance curves (Figure 2d and 2e). The latter were obtained after determining the contact point of each curve (see Materials and Methods). Force versus z-piezo and force versus tip-sample distance curves in Figure 2d and 2e are used to obtain the true sample topography, which excludes any deformation produced by the force set point used in its determination (see Materials and Methods). The true topography image of the analyzed cell is shown in Figure 2f. Comparison of the rough and true topographies (2b and 2f, note the change in color scale) illustrates the large deformations produced in the measurements (~3 μm). The thickness values extracted from the true topography were those used in the batch fitting routine to extract the Young’s modulus in every image pixel using the bottom effect-corrected analytical equation^35^. For the analysis a region of interest (ROI) around the central part of the cells was defined manually (in this case N_ROI_=189). In Figure 2g the spherical model fitted curves are shown overlaid on the experimental curves belonging to the cellular ROI. Figure 2h shows the histograms of the obtained Young’s modulus (*E*), contact point variation (Zc) and fitting accuracy (R^2^) for the analyzed cell, together with the corresponding maps as insets. The fitting routine gives the contact point variation as a fitting parameter; thus, leading to a refined true topography. Figure 2i shows the histograms of the rough topography, true topography and refined true topography as described above. As we mentioned before, the true topography exhibits higher values than the rough topography, as it accounts for the sample deformation during cell indentation. We did not observe significant variations in *E* and R^2^ by employing the true topography instead of the refined true topography. For this reason, in this work we used the true topography as a model-independent measurement of cell height. Finally, as additional output of the presented analysis, Figure 2j shows individual indexed force curves belonging to the cellular ROI with the corresponding fitted curves and fitting R^2^. More details on the procedure for force curve analysis are described in the Materials and Methods section. Data such as those shown in Figure 2 are included as additional files for all tested cells in this work, together with the raw data (see details in the Supplementary Information).

**Figure 2.**
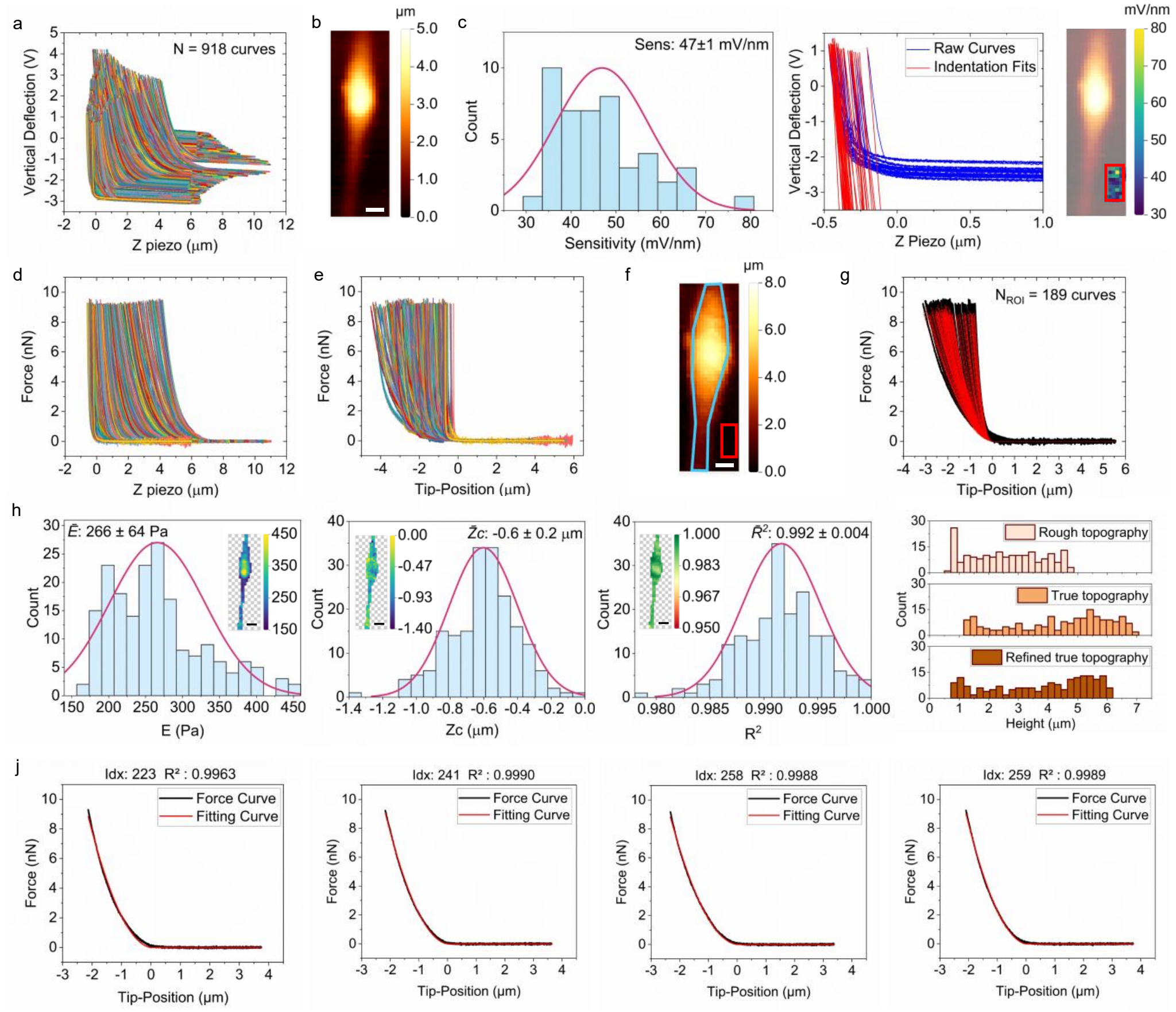
Workflow of the AFM mechanical analysis of macrophages (17×l54 force curves on an area of 28 × 90 µm^2^). (a) Raw photodetector voltage vs z-piezo curves, (b) rough topography image at the experimental force setpoint (F_set_=9 nN). Scale bar: 10 µm. (c) Optical lever sensitivity calculation (OLS=47±1 mV/nm). It includes the histogram of the slopes of the contact region of the raw curves (N_subs_=46), the fitted curves and the map of slope values superimposed on the rough topographic image, (d) calibrated force vs z-piezo curves, (e) force vs tip-sample distance curves. (f) True topography used for cell height determination and (g) results of the batch fitting showing the sphere model fitted curves (in red) overlaid on the experimental force vs tip-sample distance curves of the cellular ROI (N_ROI_=189). Experimental curves have been aligned to the refined contact point resulting from the fitting. (h) Young’s modulus, *E*, contact point variation, Zc, and R^2^ histograms resulting from the fits of the ROI force curves. The corresponding maps are shown as insets. (i) Histogram comparisons among the topographies described in the text, i.e. rough topography, true topography and refined true topography of the ROI. (j) Representative examples of force vs tip-sample distance fitted curves belonging to the cellular ROI and corresponding fitting R^2^.

It should be noted that in our analysis the force versus tip-sample distance curves spanned large indentation depths of 2-4 µm (Figure 2e). Large indentation measurements allowed full exploration of the mechanical response of macrophages to the indentation of the colloidal probe. Furthermore, they allowed an accurate *E* value to be obtained with the spherical model employed in the present study, which is more accurate than other bottom effect-corrected models, like the one for paraboloid probe geometry, at large indentations^35^.

Figures 3 and 4 show the results of the analysis for the THP-1 and BMDM-derived macrophages with typical shapes highlighted in Figure 1 (N = 22, 21 and 19 for M0, M1 and M2 from the THP-1 model, respectively; N = 18, 19 and 22 for M0, M1 and M2 from the BMDM model, respectively). To ensure a correct methodological comparison, we analyzed cellular regions exhibiting similar curvature and similar topographic variations compared to the background substrate. These regions included the cell nucleus and partially the elongated regions. Cell regions at the glass interface, where force curves were aberrant, were discarded. Extended and roughly rounded perinuclear regions typically present in the M1 phenotype from the BMDM model (see Figure 1b) were also excluded from this analysis. These areas showed a flat morphology, typical heights of around 200–500 nm and lateral extensions of tens of micrometers, thus appearing very different from the typical curvature of all the other macrophage cells. The flat morphology could affect the analysis by producing heterogeneous contact with mixed contributions from flat cell areas, curved cell areas and the glass substrate, under the micrometer-sized colloidal probe used in the experiments (R = 5 μm). Thus, only the cell nucleus area was analyzed in this case, due to the very different morphology of mouse-derived M1 macrophages compared to the shapes of the other phenotypes explored in this work.

**Figure 3.**
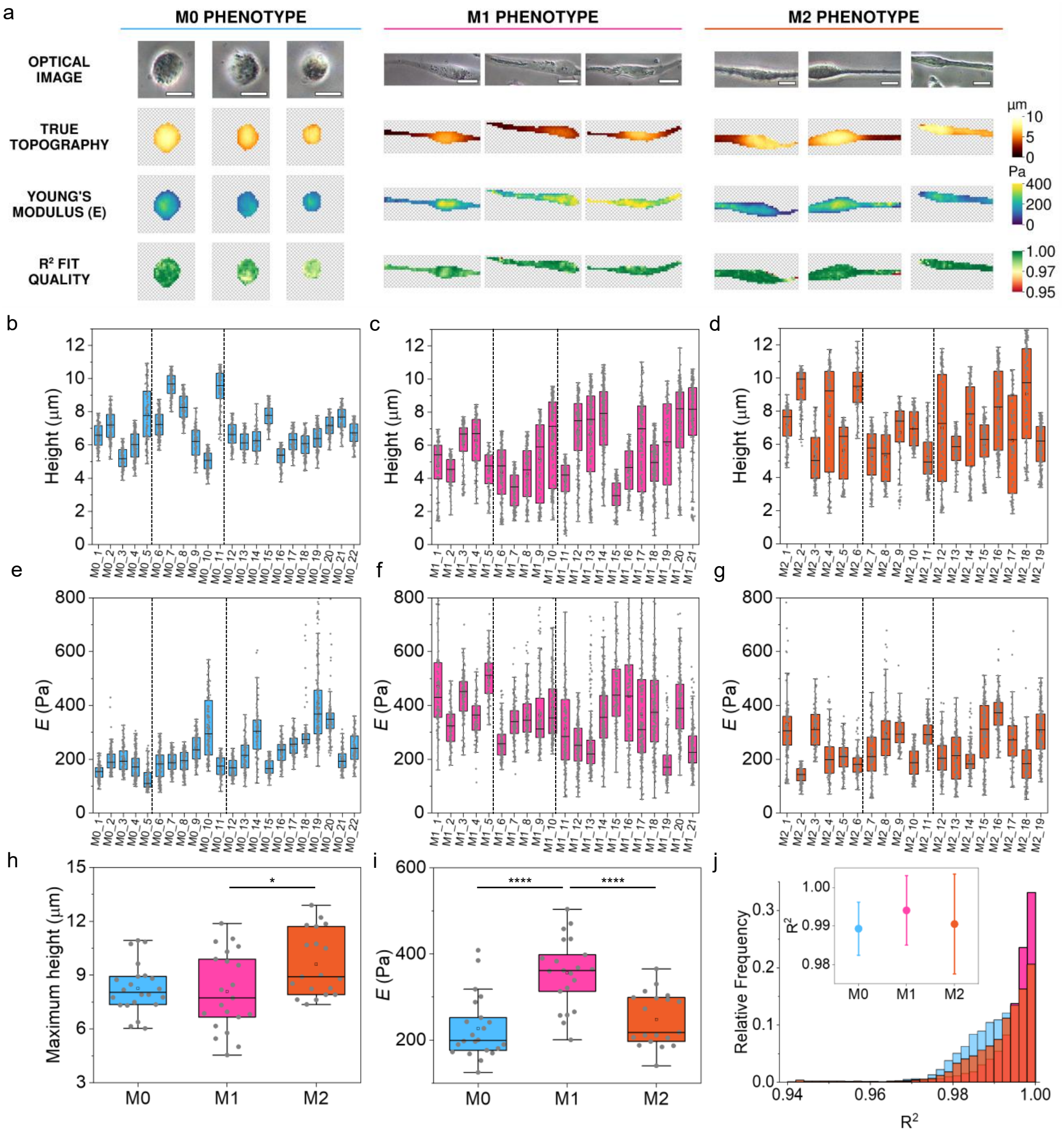
Results of the AFM analysis in the case of THP-1-derived macrophages. (a) Optical, topographic, *E* and R^2^ images of selected cells belonging to M0, M1 and M2 phenotypes. Scale bar: 10 µm (M0), 20 µm (M1, M2). (b-d) Height and (e-g) corresponding *E* of each cell ROI (N = 22, 21 and 19 respectively for M0, M1 and M2). The vertical lines in gray in (b) to (g) separate the experimental days. (h) Maximum height and (i) average *E* of every cell within each analyzed phenotype, together with the results of statistical comparison with one-way ANOVA (see Materials and Methods). (j) Normalized histograms of R^2^ values from force curves fitting with average values ± sd for each phenotype (inset).

**Figure 4.**
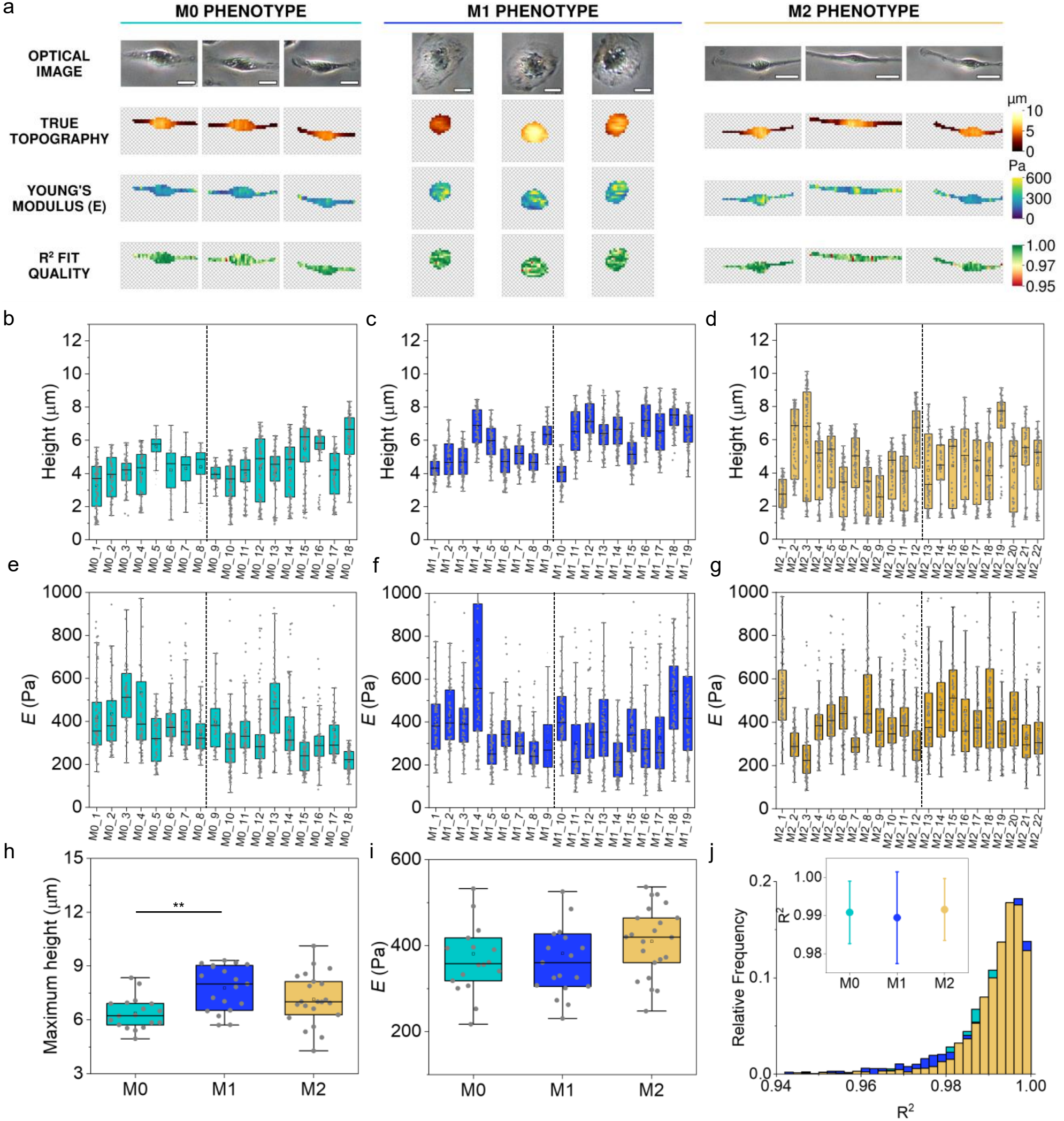
Results of the AFM analysis in the case of BMDM-derived macrophages. (a) Optical, topographic, *E* and R^2^ images of selected cells belonging to M0, M1 and M2 phenotypes. Scale bar: 10 µm for all the images. (b-d) Height and (e-g) corresponding *E* of each cell ROI (N = 18, 19 and 22 respectively for M0, M1 and M2). The vertical line in gray in (b) to (g) separates the experimental days. (h) Maximum height and (i) average *E* of every cell within each analyzed phenotype, together with the results of statistical comparison with one-way ANOVA (see Materials and Methods). (j) Normalized histogram of R^2^ values from force curves fitting with average values ± sd for each phenotype (inset).

Examples of the ROIs selected for AFM analysis are shown in Figures 3a and 4a, where the ROI true topography, *E* and R^2^ images are shown for selected cells together with the optical images collected on the same cells during the AFM experiments. At the experimental resolution (1.4 μm/pixel) the Young’s modulus maps in Figures 3a and 4a showed a contrast which is not correlated to specific cell regions, like the cell nucleus or cell elongations, exhibiting more homogeneous or more heterogeneous distributions depending on the specific cell analyzed. M0 cells in both models showed relatively more homogeneous Young’s modulus distribution compared to the other phenotypes, see the *E* images in Figures 3a and 4a. Topographic images exhibited some variations among sample types; for example, the nuclear areas of M0 and M2 cells appeared slightly higher than the nuclear areas of M1 cells in the THP-1 model. Cell height values are shown individually for each cell in Figures 3b-d and Figures 4b-d. Here, a larger spread of the height values (i.e., interquartile range) can be observed in M1 and M2 macrophages compared to M0 (THP-1), and in M0 and M2 macrophages compared to M1 (BMDM) that correlates with the increase in the projected area of the cellular ROIs (see Figures 3a and 4a). Young’s modulus values for each individual cell are shown in Figures 3e-g and 4e-g. The distribution of the Young’s modulus values was log-normal in all cases, consistent with typical findings in AFM nano and micromechanics^39–41^, see the Supplementary Figures S4a-b. The comparison between phenotypes is reported in Figures 3h-i and Figures 4h-i, using the maximum topographic height and the average *E* calculated for each cell as metrics. Significant differences among cell types were observed regarding the maximum height, see the comparison between M2 and M1 in the THP-1 model (Figure 3h, p<0.05) and between M0 and M1 in the BMDM model (Figure 4h, p<0.01). However, the most remarkable result was obtained for the Young’s modulus in the THP-1 case, where we found strong statistical evidence that the cellular *E* is higher after polarization to the M1 phenotype (p<0.0001), while M0 and M2 phenotypes do not differ significantly (Figure 3i). This significantly higher *E* obtained in the case of the M1 phenotype from THP-1 is remarkable also considering the limited cell datasets that can be explored by AFM. The higher E values found in M1 cells are consistent with previous findings that employed other models, like the mouse-derived cell line RAW 264.7 and human-derived macrophages^26,29,30^. In the case of the BMDM model, instead, polarization into M1 and M2 did not produce any significant change in the cellular *E* (see Figure 4i). Furthermore, in this case, M0 and M2 macrophages showed higher *E* compared to the corresponding cell types in the THP-1 model, with similar Young’s modulus values in M1 cells in the two models.

Finally, for both cell models, the application of the sphere model with bottom effect correction used to analyze force curves at high indentations gave very good fitting accuracy (R^2^ ≈ 0.991 on average, see Figures 3j and 4j). We also note the reproducibility of the results, which were collected on different days and with different cell batches (separated in the figures by the vertical lines).

To obtain comprehensive information, we also analyzed macrophages with other shapes beyond the typical ones associated with each phenotype in both models; results are shown in Figures 5 and 6 for the THP-1 and BMDM cases, respectively. In THP-1 samples, small cells with rounded shapes were found in all samples, showing very bright optical contrast and poor adhesion on the glass-bottom Petri dishes, a fact that made their characterization by AFM not possible. We attributed these characteristics to undifferentiated monocytes^6^, which are primarily circulatory cells with poor adherence before differentiation into tissue-resident macrophages^34^. Some of these cells are visible outside the marked areas in Figure 5a-5c. Adherent cells with circular shapes (M0 and M1 samples) or slightly elongated shapes (all samples) were found (see the cells highlighted in blue and light orange in Figure 5a-c). However, they did not show significant Young’s modulus differences between phenotypes, see the comparison with Welch’s t-test for circular cells (M0 vs M1; Figure 5d and 5g) and one-way ANOVA for spindle-like cells (Figure 5e and 5h), a fact that supports their identification as unpolarized M0 macrophages from the mechanical point of view^5,6^. Furthermore, branched or dendritic cells were occasionally found in M2 samples and they were also characterized by AFM, showing very homogeneous *E* profiles and non-significant differences with the elongated M2 cells (Figure 5f and 5i), see also the mechanical images associated with these cells, provided as additional files (see Supplementary Information). Figure 6a-c shows that the typical shapes of M0, M1 and M2 macrophages were ubiquitous in BMDM-derived samples. However, Young’s modulus values of all mouse-derived cells were in a similar range (see Figure 6d-f) and no statistical differences were found either comparing a specific cell shape with all the other shapes within the same phenotype or comparing the same cell shape among different phenotypes (see Figure 6g-i). Some topographic differences were found in cells with different shapes for the THP-1 and the BMDM type (see Supplementary Figures S5 and S6).

**Figure 5.**
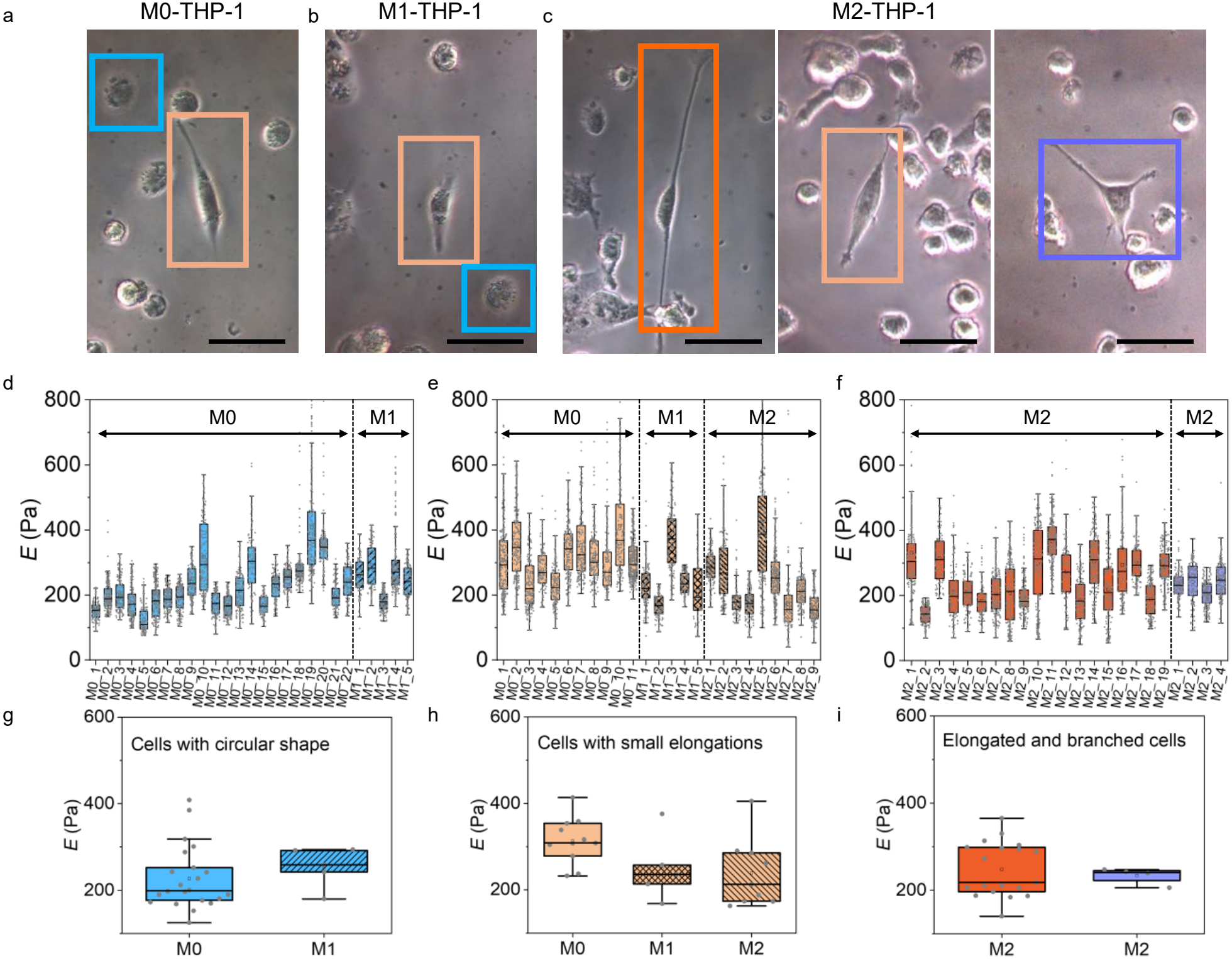
Optical images showing other shapes found in M0 (a), M1 (b) and M2 (c) samples from THP-1 beyond characteristic phenotypical shapes. Scale bar: 50 µm for all the images. (d-e) Young’s modulus values associated with well-adhered cells of circular shape (d) and spindle-like cells (e) present in the different samples. (f) Young’s modulus values of elongated cells (in orange) and branched, roughly triangular cells (in violet), present in the M2 samples. Vertical lines in gray separate different experimental days. (g–i) Average *E* for each of the analyzed shapes. Statistical analysis, i.e. Welch’s t-test (g, i) and one-way ANOVA (h), gave non-significant differences between phenotypes in all cases. The only significant difference was found between circular adhered cells and cells with small elongations in M0 samples, the latter showing higher *E* (Welch’s t-test, p<0.001).

**Figure 6.**
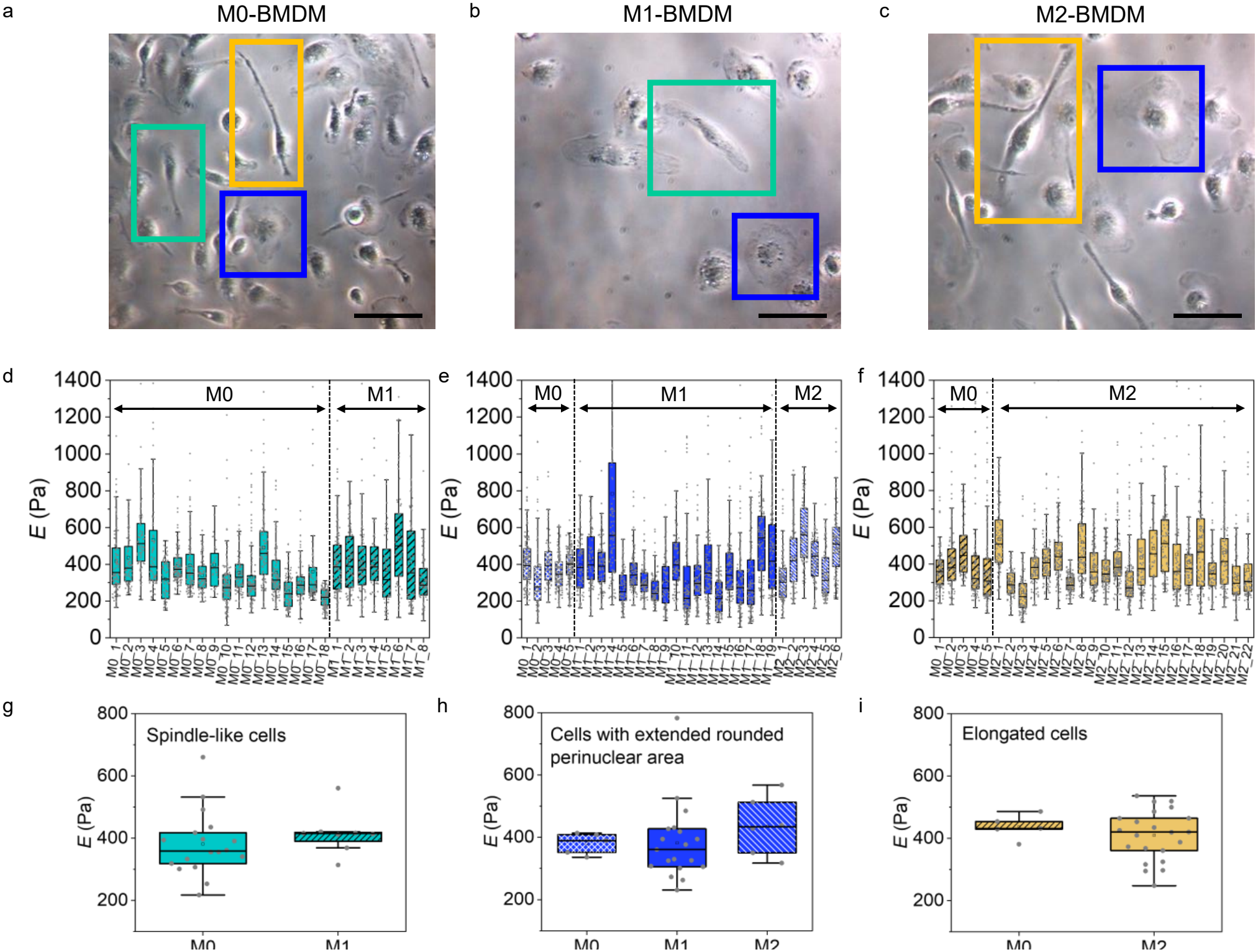
Optical images showing other shapes found in M0 (a), M1 (b) and M2 (c) samples from BMDM beyond characteristic phenotypical shapes. Scale bar: 50 µm for all the images. (d–f) Young’s modulus values associated with spindle-like cells (d), cells with large, roughly rounded perinuclear area (e) and elongated cells (f) present in the different samples. The gray vertical line in (d)–(f) separates the different experimental days. (g-i) Average *E* for each of the analyzed shapes and results of the statistical analysis, i.e. Welch’s t-test (g, i) and one-way ANOVA (h), giving non-significant differences between group means in all cases.

The outputs of the analysis of the cells in Figures 5 and 6, with all *E*, true topography and R^2^ maps are provided as additional files, see Supplementary Information.

## 3. Discussion

A comprehensive mechanical characterization of macrophage cell types from two different cell models (THP-1 and BMDM) has been reported by means of large indentation force volume AFM measurements with colloidal probes. On the methodological side, we highlight that we used the recently derived sphere model with bottom effect correction^35^. This model considers a spherical probe indenting a laterally infinite, homogeneous and elastic thin film bonded to a hard substrate, without considering adhesive and viscoelastic effects^35^. At high indentations (*δ*), the model predicts a force vs tip-position curve that diverges from other models used in the literature, e.g. the Sneddon model for a spherical indenter without bottom effect correction (BEC)^42^ or the paraboloid model with BEC and without it (Hertz’s model). This divergence between models leads to different *E* values from curve fitting depending on the model used.

Specifically, in the high indentation regime (*δ*/R>0.4), the sphere model without BEC (Sneddon’s model) overestimates the Young’s modulus due to the bottom effect, while the paraboloid model with BEC underestimates *E*^35^. This effect is shown in Figure S7 where the same M0 cell from THP-1 was analyzed using the three models, leading to average values *E*_Sneddon_ = 428±73 Pa, *E*_Parab BEC_ = 156±53 Pa and *E*_Sphere BEC_ = 193±56 Pa. Deviations of +122% and −19% are obtained for Sneddon’s and paraboloid with BEC models with respect to the sphere model used in the present work. The choice of the correct geometry and model for data analysis is expected to translate into more physically meaningful values of *E*, thus benefiting the standardization of AFM measurements^43,44^.

A second methodological aspect to highlight is that we considered high indentation measurements. To achieve the highest possible indentation (*δ*) of macrophages, the applied force during the experiments was set to 8-9 nN (see Materials and Methods) and *δ* values in the range 2-4 μm were obtained in areas close to and including the cell nucleus (see Figures 2 and S7). For these high indentation measurements our results gave average *E*_M0_ = 227±73 Pa, *E*_M1_ = 356±81 Pa and *E*_M2_ = 248±62 Pa for THP-1-derived macrophages and average *E*_M0_ = 381±104 Pa, *E*_M1_ = 382±124 Pa and *E*_M2_ = 410±81 Pa for BMDM, with very good fitting accuracy using a single fitting function to the high indentation data (see Figures 2j, 3j, 4j and S7, and insets). In this respect, we note that other works have reported separate fitting of the low and high indentation part in the force curves^26,29,30^.

The obtained *E* values, of the order of a few hundred pascals, confirm the very soft nature of macrophage cells. To put our results in the context of the current literature, we observe that mechanical analysis of macrophages by AFM has been performed using different conditions in terms of probe geometry^22^, the indentation depth during measurements, the indentation depth used for force curves analysis, the model used for the analysis^29^ and the cell sample preparation. Experiments with nanometric probes and fixed cells^14,21,23,24,34,45^ are very far from the experimental conditions explored in this work and can give very different results; thus, they are not considered here. A more direct comparison considers experiments performed with colloidal spherical probes and living cells. Galluzzi and coworkers analyzed mechanical characteristics of living macrophages from different cell models (THP-1^25^, mouse-derived RAW 264.7 line^25–28^ and human-derived primary macrophages^30^) using probes similar to those employed in this work (probe radius R = 5 µm and cantilever k=0.2 N/m), lower indentations (*δ*<2 µm) and an analytical model of a paraboloid indenter with BEC that assumes the cell is weakly bonded to the substrate^27^. The approach velocity used in those works was similar to ours, a fact that facilitates the comparison, given the viscoelastic nature of cells and the sensitivity of AFM indentation experiments to the indentation speed^46,47^. The prediction of the employed model in these references^25–27,30^ is a higher *E* compared to the bottom bonded model for the same geometry. Under this assumption, small differences among phenotypes, i.e. E_M1_ > E_M2_ > E_M0_ in the three analyzed models have been reported, with absolute Young’s modulus values slightly lower than those found in this work (E_M0_~180 Pa, E_M1_~270 Pa for THP-1^25^; E_M0_~160 Pa, E_M1_~220 Pa, E_M2_~200 Pa, for human macrophages^30^). For mouse BMDM a different tendency among phenotypes has been reported, with E_M2_>E_M1_>E_M0_ in force spectroscopy experiments conducted with colloidal probes^31^. In our experimental conditions, the mechanical behavior of THP-1-derived macrophages aligns with the above-mentioned reports, while about 1.5 times higher *E* values were found in the BMDM model compared to the basal *E* level in M0 macrophages from THP-1, with no significant differences among the phenotypes (Figure 4i).

We hypothesized that the non-significant phenotypic differences found in the BMDM model could be attributed to the restricted choice of the cellular ROI in the case of the M1 phenotype (see Figure 4a). For this reason, we performed additional mechanical analysis, including the image pixels belonging to the perinuclear regions in BMDM-derived M1 cells. Results are shown in Supplementary Figure S8. In this case, while we observed a slight increase in the *E* values in all analyzed cells (Figure S8f and S8g), we still did not find significant differences between the three phenotypes’ average *E* (Figure S8g). The cellular shape typical of M1 cells in the BMDM samples and characterized by extended flat perinuclear areas (thickness = 200-500 nm) was markedly different from the typical shapes assigned to the other macrophage phenotypes, which had the common denominator of a rather curved cellular profile. Due to the limited thickness of these areas, inaccuracies in contact point determination can strongly affect the true sample height and the extracted Young’s modulus values. For these cells, a lower average R^2^ = 0.975 ± 0.033 was also obtained (see Figure S8h and inset). Furthermore, from the biological perspective, regions of M1 macrophages with a high content of fibrous proteins (F-actin, α-tubulin), possibly giving rise to an increase in *E*, have been shown not to be limited to peripheral cell areas^15,16,27^. Thus, we considered restricting the analysis to the nuclear region as the best methodological approach from the AFM perspective, as reported in Figure 4a. Morphometric analysis was done considering the entire cell area of these macrophages, i.e. including flat perinuclear regions (see the cellular ROIs for both analyses included as supplementary files).

To explain the different results among cell models (Figure 3 and 4), we observe that the higher Young’s modulus of M1 compared to M0 and M2 in THP-1 is likely related to the signaling triggered by the M1 stimuli (LPS and IFN-γ). Stimulation of macrophages with LPS and IFN-γ induces a myosin II-dependent contraction followed by actin-dependent cell spreading^48^. In primary human macrophages, it has been shown that IFN-γ increases cell stiffness while LPS alone does not, and stiffness correlates closely with F-actin content^49^. The M1 stiffening observed here may therefore be driven mainly by IFN-γ, which was used at a ten-fold higher concentration than LPS (25 vs 2.5 ng/mL, see the Materials and Methods section). THP-1 cells are also known to respond less to M2 stimulation than primary macrophages. For example, they fail to express the M2 surface marker CD163 after IL-4/IL-13 treatment, whereas their response to M1 stimulation is closer to that of primary macrophages^50^. This may explain why M2 cells remain mechanically similar to M0 cells.

In the BMDM model, polarization did not change the Young’s modulus, and all three phenotypes showed values similar to those of M1 cells in the THP-1 model. We propose that this reflects the different differentiation routes of the two models. BMDM are generated by seven days of culture in M-CSF (Materials and Methods), and activation of the M-CSF receptor drives actin polymerization, cell spreading and adhesion in macrophages^51^. THP-1 cells, in contrast, were exposed to PMA for 48 hours without a resting phase. Under these conditions THP-1 cells can be less mature than bone marrow-derived macrophages, with smaller cytoplasm and weaker adhesion (the smaller cytoplasm was confirmed optically, see the phase contrast images in Figure 1). It has been shown that a resting period after PMA removal makes them more like primary macrophages^52,53^. The difference observed in the two models may therefore depend mainly on the duration and completeness of differentiation^53^, rather than on a weaker effect of PMA on the cytoskeleton. BMDM may reach the polarization step with a high level of polymerized actin and higher stiffness, leaving limited room for further stiffening upon LPS/IFN-γ stimulation. In this view, LPS/IFN-γ stimulation brings THP-1 cells to a cytoskeletal state that BMDM cells already reach during M-CSF differentiation.

A final comment concerns the presence of additional contributors to the mechanical differences observed in both models. Figure 7a summarizes the results obtained in this work, comparing statistically the same phenotype between the two models. In our experiments we analyzed parameters of the cellular shape without other functional analysis of fluorescent biomarkers, which can be correlated with the amount and distribution of fibrous proteins of the cell cytoskeleton, e.g., F-actin^16,25,30^. Our results showed an apparent lack of correlation between the cellular shape and the Young’s modulus. For example, in THP-1 macrophages, the higher Young’s modulus observed for the M1 phenotype compared to the M2 phenotype cannot be correlated with cell elongation, which is a common feature of M1 and M2 phenotypes (see Figure 1a and 1e-h). Similarly, in BMDM-derived macrophages, different shapes, degree of elongation and circularity in M0, M1 and M2 (Figure 1i-l) did not translate into distinct *E* values. To find a morphological parameter that scales with *E*, we observed that THP-1 and BMDM macrophages exhibited visually different topographic heights (see Figures 3h, 4h and S3). Thus, we propose the maximum height divided by the projected area as a metric of the cellular aspect ratio that can also affect macrophage elastic behavior. Results are shown in Figure 7b and 7c. Significant differences in the aspect ratio were only found between the M0 and M2 phenotypes of the explored models, consistent with the *E* results (Figure 7b), with lower aspect ratios in M0 and M2 phenotypes from the BMDM model. In Figure 7c, *E* vs cellular aspect ratio is shown for all the tested cells in this work (179 cells, including all different shapes present in samples). Correlation analysis gave a statistically significant, moderate inverse relationship between the Young’s modulus and the aspect ratio (Pearson’s coefficient r=−0.42, p<0.0001). Similar results were obtained using the cell volume as a metric, see Supplementary Figure S9. This tendency is consistent with spreading models of actin polymerization, increased membrane tension upon cell adherence on a substrate and crowding due to volume reduction^54,55^. Thus, morphometric cell parameters (flat morphology, spreading area, volume), rather than a specific cellular shape, also affect the macrophages’ Young’s modulus in addition to phenotypic polarization.

**Figure 7.**
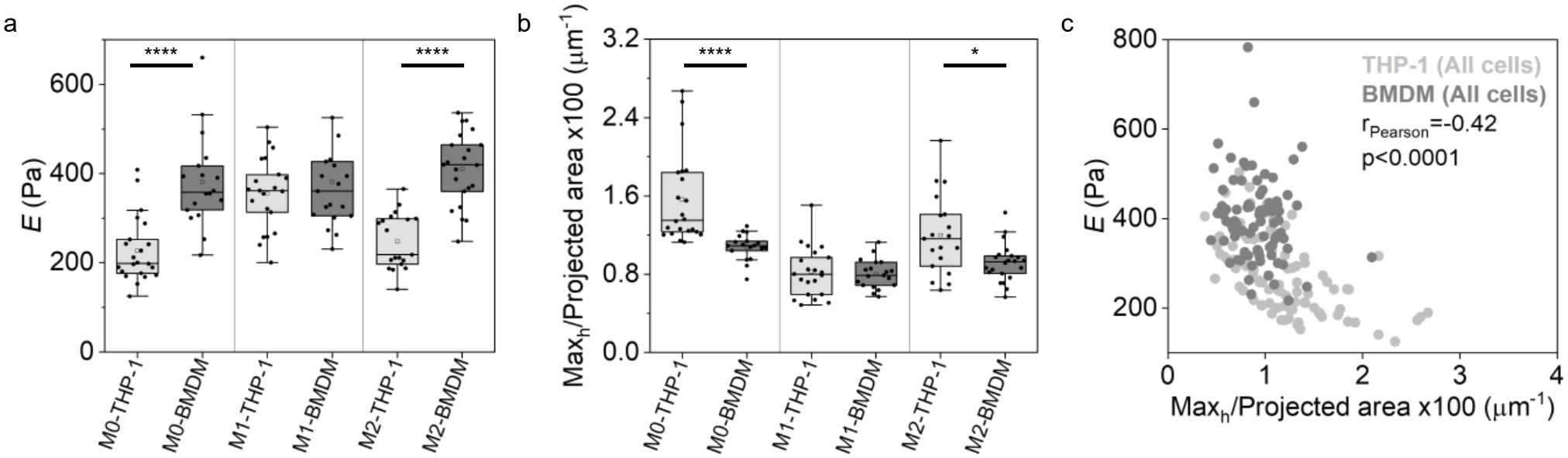
(a) Young’s modulus values highlighting the differences between the THP-1 and BMDM cell models. Vertical lines separate the M0, M1 and M2 phenotypes. (b) Corresponding plot of the aspect ratio, i.e. the maximum height divided by the cell projected area. In both graphs the results of comparison with Welch’s t-test are included for statistically different samples. (c) *E* vs cellular aspect ratio with the results of the correlative analysis considering all cells from THP-1 and BMDM as a single group and all shapes.

### 4. Conclusions

In conclusion, we present the results of the large indentation mechanical analysis by AFM of M0, M1 and M2 macrophage phenotypes from two recognized cell models, i.e. THP-1 and BMDM. Our methodology is based on a newly developed analytical equation for spherical probes that accounts for the bottom contact effect to obtain quantitative values of the Young’s modulus in experiments performed with colloidal probes at large indentations. Colloidal probes are being extensively used for cell mechanical analysis, guaranteeing soft contact, low risk of cell damage and reduced contamination of the tip. Our approach is useful to analyze force curve-based AFM data in experiments aimed at testing many cells with low resolution maps (tens to hundreds of cells, ~600–1400 force curves/image, ~1–3 µm^2^/pixel), i.e. for quantitative mechanical cell phenotyping at the level of cell populations. Also, our methodology can benefit data reproducibility and methodological standardization, which are important needs of AFM research in the field of cells and soft matter characterization. Our results show low values of the Young’s modulus, in the range 200-400 Pa, for all the tested cells and significantly higher *E* values for M1 macrophages only in the case of the THP-1 model, while all phenotypes exhibited similar *E* close to 400 Pa in the BMDM model. Macrophage cells show an intrinsically rich morphology; furthermore, they undergo important morphological changes upon polarization. We found that these morphological transformations alone cannot explain the mechanical differences observed by AFM in the two models. A possible contribution to the observed Young’s modulus beyond phenotypic polarization can be found in the cellular aspect ratio, which affects different regulatory processes, such as mechanotransduction and adhesion in many cell types, including macrophages. Finally, our results show a different mechanical behavior between the cell line and primary mouse-derived macrophages, a fact that confirms the lack of uniform behavior among the different cell models, also due to the distinct differentiation protocols, that could be an additional source of discrepancies in *in vitro* experiments on host– pathogen interactions, immunology and drug development for immunotherapy.

## Materials and Methods

### Cell Culture and Differentiation

#### THP-1 Cell Line

THP-1 cells (human acute monocytic leukemia cell line, ATCC TIB-202) were maintained in suspension in T-75 flasks in RPMI-1640 medium (Gibco, Thermo Fisher, Ref. 11835063) supplemented with 10% fetal bovine serum (FBS; Life Technologies, Ref. 10270106), 1% penicillin-streptomycin (P/S; Gibco, Thermo Fisher, Ref. 15140122), and 0.05 mM β-mercaptoethanol (BME; Gibco, Thermo Fisher, Ref. 31350010), at 37 °C in a humidified atmosphere with 5% CO2. All percentages are expressed as v/v%. Medium was replaced every 2–3 days, and cells were passaged weekly to maintain a density between 2 × 10^5^ and 1 × 10^6^ cells/mL. To avoid phenotypic drift, only cells between passage 6 and 15 were used in this work, and cultures were discarded after passage 20. Cells were routinely tested for mycoplasma contamination. For differentiation, cells were seeded at 5 × 10^5^ cells/dish onto FluoroDish glass-bottom culture dishes (35 mm diameter; World Precision Instruments) in the culture medium described above. Cells were first differentiated into unpolarized macrophages (M0) by incubation with 20 ng/mL phorbol 12-myristate 13-acetate (PMA; Sigma-Aldrich, Ref. P1585) for 48 h. From this M0 population, two independent polarization branches were established: (i) polarization toward the pro-inflammatory M1 phenotype, by co-incubation with 25 ng/mL human interferon-gamma (hIFN-γ; GenScript, Ref. Z02915) and 2.5 ng/mL lipopolysaccharide from *Escherichia coli* (LPS; Sigma-Aldrich, Ref. L2630) for 24 h; and (ii) polarization toward the anti-inflammatory M2 phenotype, by stimulation with 100 ng/mL human interleukin-13 (hIL-13; BioLegend, Ref. 571002) and 100 ng/mL human interleukin-4 (hIL-4; BioLegend, Ref. 574302) for 72 h.

#### Mouse Bone Marrow-Derived Macrophages (BMDMs)

All animal procedures were approved by the CEEA-PCB (Barcelona), under protocol number 24-014. Bone marrow was isolated from the femur and tibia of 6-week-old C57BL/6 male and female mice (day 0) following euthanasia by cervical dislocation. Red blood cells were lysed using RBC lysis buffer (Invitrogen, Ref. 00-4333-57) for 5 min.

Bone marrow cells were plated in p100 Petri dishes (Sarstedt, Ref. 821473001) in 8 mL of R10 medium supplemented with 30% macrophage colony-stimulating factor (M-CSF)-conditioned L929 supernatant (M-CSF/L929) and cultured for 7 days to induce differentiation into BMDMs. R10 medium consisted of RPMI-1640 (Gibco, Ref. 11835030) supplemented with 10% heat-inactivated FBS (Gibco, Ref. A5256801), 1% P/S, 0.1 mM non-essential amino acids (NEAA; Thermo Scientific, Ref. 11140050), 1 mM sodium pyruvate (Gibco, Ref. 11530396), 1 mM HEPES (Thermo Scientific, Ref. 1447837), and 0.05 mM BME (Gibco, Ref. 31350010). M-CSF/L929 conditioned supernatant was generated by culturing L929 cells in R10 medium lacking HEPES and BME for 15 days and stored at –80 °C until use. Twenty plates were prepared per animal. On day 3, each plate received 3 mL of fresh R10 + 30% M-CSF/L929. On day 7, adherent BMDMs were harvested using cell scrapers and reseeded at 5 × 10^5^ cells/mL onto FluoroDish glass-bottom culture dishes (35 mm diameter; World Precision Instruments) in R10 medium without BME and containing the appropriate stimulants. Cells were incubated for 24 h under the following conditions/stimulants: M0 macrophages were maintained in R10 medium alone; M1 macrophages were polarized with 100 ng/mL lipopolysaccharide (LPS; InvivoGen, Ref. tlrl-eklps) and 50 ng/mL recombinant mouse interferon-gamma (mIFN-γ; BD Biosciences, Ref. 554587); M2 macrophages were polarized with 20 ng/mL recombinant mouse interleukin-4 (mIL-4; PeproTech, Ref. 214-14).

### Macrophage analysis

#### Quantitative polymerase chain reaction (qPCR)

Samples consisted of 1×110^6^ to 2×l10^6^ cell pellets, washed 3 times with PBS to remove culture medium. Each cell pellet was resuspended in 300 µL of lysis buffer present in the Monarch® Total RNA Miniprep Kit (New England Biolabs, Ref. T2010S) at room temperature under RNase-free conditions. RNA extraction was completed following the manufacturer’s instructions, and RNA was kept on ice thereafter. RNA purity and concentration (ng/µL) were assessed on a NanoDrop 8000 spectrophotometer (Thermo Scientific). Reverse transcription (RT) was performed with the ProtoScript® II First Strand complementary DNA (cDNA) Synthesis Kit (New England Biolabs, Ref. E6560S) following the manufacturer’s instructions. A control reaction was included where the RT enzyme was omitted. The obtained cDNA was diluted 1:10 and used as a template for real-time amplification in qPCR.

For THP-1-derived samples, Mix 1 was prepared with 2 µL of diluted cDNA (4 ng total cDNA) and 6 µL of nuclease-free water. Mix 2 included 1 µL each of 10 µM forward and reverse primers and 10 µL of Power SYBR™ Green PCR Master Mix (Applied Biosystems). Human primer sequences are listed in Table I. Each primer pair was tested with a control reaction consisting of water instead of cDNA. Mix 1 and Mix 2 (20 µL final volume) were combined in a MicroAmp™ Fast Optical 96-Well Reaction Plate (Applied Biosystems, Ref. 4346906), sealed with MicroAmp™ Optical Adhesive Film (Applied Biosystems, Ref. 4360954), and run on a StepOnePlus™ Real-Time PCR System (Applied Biosystems). The thermocycler program consisted of an initial denaturation at 95 °C for 10 min and amplification for 40 cycles of 15 s at 95 °C and 1 min at 60 °C. A melt curve was acquired to verify amplification specificity, with an extra step at 95 °C for 15 s, 60 °C for 1 min and 95 °C for 15 s. Relative fold increase in gene expression was calculated with the 2^(−ΔΔCt) method^56^ and normalized to the endogenous gene GAPDH. For BMDMs, reaction preparation was automated using the QIAgility pipettor (Qiagen). Total reaction volumes were halved, with 0.2 mL PCR tubes (Labbox, Ref. PCRP-002-1K0) containing 1 μL of cDNA (8 ng total cDNA) and proportionally adjusted volumes for other components. Mouse primer sequences are listed in Table II. Sealed tubes were run on a Rotor-Gene Q thermocycler (Qiagen), following the same thermocycling program as for THP-1 samples. In this case, gene expression was calculated using the 2^(−ΔΔCt)^56^ method, normalized to the endogenous gene β-actin (selected as the most stable of three candidate reference genes, GAPDH, HPRT and β-actin, across stimulation conditions).

**TABLE I.**
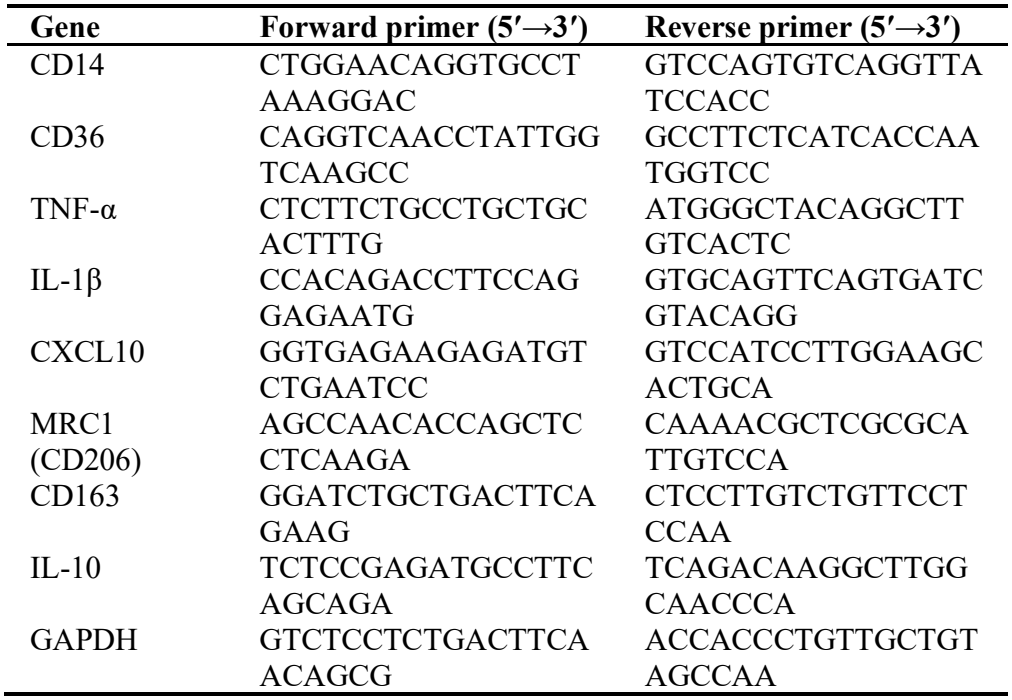
Human oligonucleotide primers used for qPCR.

**TABLE II.**
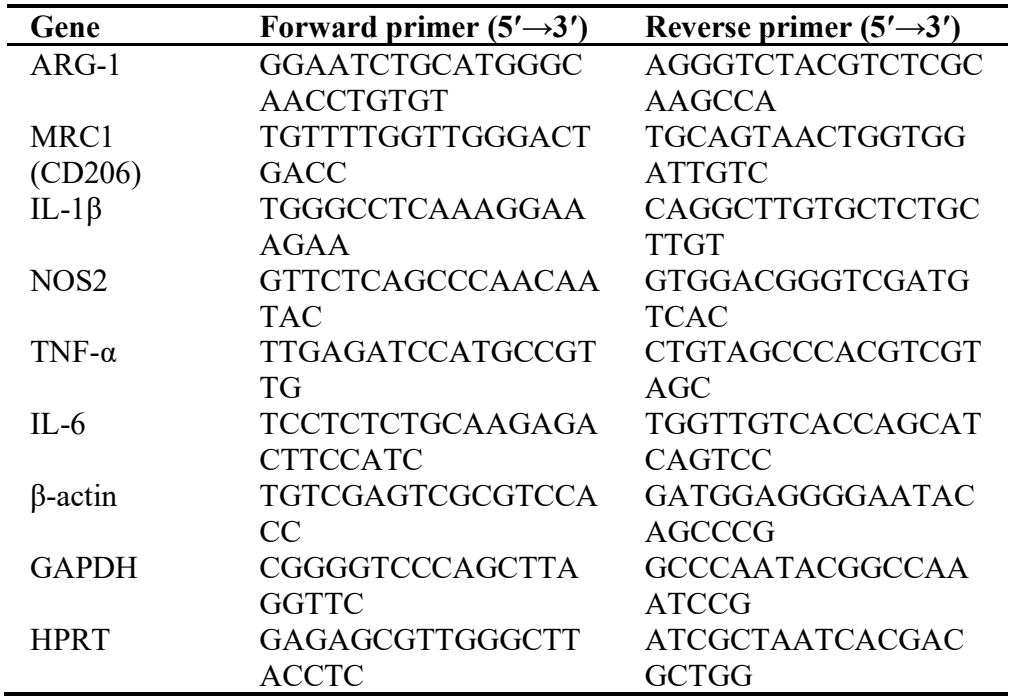
Mouse oligonucleotide primers used for qPCR.

#### Flow cytometry

M0 cells from BMDMs were seeded in 12-well plates at 5×10^5^ cells/well and cultured in R10 medium. Polarization was performed under previously optimized conditions: 100 ng/mL LPS + 50 ng/mL IFN-γ for M1 macrophages and 20 ng/mL IL-4 for M2 macrophages for 24 hours. The immunostaining protocol was performed entirely with ice-cold solutions, with cells maintained on ice to preserve viability. Cells were washed twice with cold HBSS (ThermoFisher Scientific, Ref. 14175095), gently scraped, and transferred into 1.5 mL Eppendorf tubes. They were centrifuged at 1800 rpm for 5 minutes at 4 °C using an Eppendorf 5425 R microcentrifuge. To block non-specific Fc-mediated interactions, the cell pellets were resuspended in 100 μL of 2% (w/v) BSA (Merck, Ref. A7638) in PBS containing Mouse BD Fc Block™ (BD Biosciences, Ref. 553142; 1:100). After a 10-minute incubation, Alexa Fluor® 647 anti-mouse CD206 (BioLegend, Ref. 141712; 1:100) and Alexa Fluor® 488 anti-mouse I-A/I-E (MHC-II) (BioLegend, Ref. 107616; 1:200), or the matched isotype controls AF647 Rat IgG2a, κ (BioLegend, Ref. 400526) and AF488 Rat IgG2b, κ (BioLegend, Ref. 400625) at the same dilutions, were directly added and incubated for 30 minutes on a rotating plate. Cells were washed twice with the BSA solution and resuspended in 250 μL PBS. For viability assessment, 1 μL of propidium iodide (Sigma-Aldrich, Ref. 81845; 0.1 mg/mL in PBS) was added to each tube. Samples were immediately analyzed using a spectral flow cytometer (Cytek® Aurora).

#### ELISA

BMDMs were seeded in 6-well plates at 1×10^6^ cells/well (0.5×10^6^ cells/mL, 2 mL/well) in R10 medium and stimulated for 24 hours under the conditions shown in Fig. S2a (C0, R10 only; C1 and C3, mouse IL-4 at 10 and 20 ng/mL; C5–C8, LPS + mouse IFN-γ at 100 + 50, 100 + 25, 50 + 50 and 50 + 25 ng/mL). 1.5 mL of culture medium was collected from each condition, centrifuged at 4 °C for 5 minutes at 2800 rpm, and supernatant was stored at –80 °C to quantify TNF-α release by sandwich ELISA using the Mouse TNF-α DuoSet ELISA kit (R&D Systems, Bio-Techne, Ref. DY410-05). Plates were pre-coated with the capture antibody at 4 °C overnight, the day before use, following the kit instructions, and TNF-α concentrations were interpolated from the standard curve.

### Atomic Force Microscopy (AFM)

Measurements were performed using a JPK NanoWizard IV atomic force microscope (Bruker) operated in Quantitative Imaging (QI) mode, which allows simultaneous acquisition of topographical and mechanical information at each pixel through force– distance curves. All experiments were conducted in cell culture medium at 37 °C, using a heated sample stage to preserve physiological conditions throughout data acquisition.

Spherical polystyrene colloidal probes (R = 5 µm, Novascan) were employed, with a nominal spring constant of 0.12 N/m. The actual spring constant of each cantilever was confirmed to be the same as the nominal value prior to each experiment using the thermal noise method.

Force curves were acquired with the following conditions: Z-range = 8 μm, force setpoint = 8-9 nN, approach/retraction velocity = 40 μm/s and sampling rate = 40 kHz. This setpoint was selected to ensure sufficient indentation depth for accurate mechanical characterization with the spherical model used in this work. Scan area was adjusted for each cell according to its size and morphology, with a pixel size of approximately 1.4 µm (median pixel area 1.9 µm^2^/px across all measurements. Round cells, like those of the M0 phenotype in THP-1 and the M1 phenotype in BMDM, were typically scanned over square areas of approximately 35×35–40×40 µm^2^, whereas elongated cells (M1/M2 in THP-1; M0/M2 in BMDM) were scanned over rectangular areas of approximately 30×50–30×90 µm^2^ to adequately capture their morphology. Force–curve datasets ranged from approximately 625 to 1400 curves/map. Under these conditions, acquisition time was ~7 min/map.

### Data analysis

Each dataset belonging to a single cell was processed and analyzed with a custom-made software tool referred to as Mechanotool, which will be described in detail in a separate publication (manuscript in preparation) to quantify the sample’s mechanical properties. First, a rough topography image was generated with the minimum Z-piezo values of each curve corresponding to the experimental force set point (the minimum Z-piezo for the substrate was set to 1 nm). The rough topography was then used to segment the dataset into the bare substrate and the region of interest (ROI). Segmentation was done manually using a polygonal selection tool to exclude low-quality curves, e.g. those at the cell-glass boundary.

Using the selected substrate region, the optical lever sensitivity (OLS) was obtained from a linear fit of the cantilever deflection contact region of each curve (units in mV/nm). The final OLS was defined as the mean value from a Gaussian fit to the distribution of these slope values. For each curve, a separate linear fit of the non-contact baseline segment was also performed. In both substrate and ROI cases, the optimal segment of the curve to be fitted was determined automatically using a sliding-window algorithm^57^. The endpoint of the baseline fit was chosen as the contact point (Z_C_), and the line given by this fit was subtracted from each curve to correct for the baseline offset and tilt.

Force curves were calibrated using the cantilever’s nominal spring constant value and the OLS. The contact point identified previously was applied, and the force-indentation curves were computed by subtracting the vertical deflection from the Z-piezo variation.

A low-force set-point topography image was generated by applying a predefined setpoint to the force vs. z-piezo curves. This setpoint was set to five times the standard deviation of the mean baseline noise. The true topography image was then produced by correcting the height values for the indentation depth reached at the applied force setpoint, removing the topographic variation introduced by probe–sample indentation.

Finally, the force-indentation curves of the ROI were fitted using the spherical indenter model with bottom effect correction^35^, with the Young’s modulus (*E*) and the contact point variation (Z_C_) left as free parameters. The fitted parameters were mapped across the ROI for visualization and displayed alongside the corresponding R^2^ maps, allowing the fitted values and quality of the fits to be visually inspected. The fitted Z_C_ can optionally be used to generate a subsequent sample height (refined true topography), using it as input in a second batch fitting routine. As the refined true topography did not significantly affect the *E* results, only the results of the first batch fitting routine are shown in this work. Additional repository files contain all analysis outputs and original QI data for each cell (see Supplementary Information). True topographic data are reported in this work. Cell topographic profiles were found to be smooth in the tested cells, without spikes. Thus, the highest pixel value from true topography images of each cell is reported as the maximum height in Figures 3h, 4h, S5d–f and S6d–f, considering this pixel height as a spatially integrated measurement over the probe-sample contact area.

### Morphometric Analysis

Using the true topographic images obtained by AFM, cells were segmented and several morphometric parameters were calculated. For each topographic image, the substrate plane was corrected by subtracting the median height of the pixels outside the cell area, setting it to Z = 0 nm. Cells were then segmented by automatically thresholding the height map (Otsu’s method, on the image normalized to [0, 1]^58^) and applying marker-based watershed segmentation; small objects were removed and internal holes were filled using morphological operations^58,59^. When adjacent cells were occasionally not properly separated, a manual line was drawn on the height image, and the mask was split along this line using a Bresenham algorithm^60^, with each resulting region analyzed independently.

All metrics were computed using custom Python software and the scikit-image library^59^. Pixel size (*Δx*, µm/pixel) was derived from the scan size and pixel resolution metadata of each file.

#### Cell projected area

The projected area (*A*) was calculated as the number of pixels belonging to the segmented cell mask multiplied by the physical area of one pixel (µm^2^):

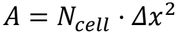

where *N*_cell_ is the number of pixels within the binary cell mask and *Δx* is the pixel size in µm.

#### Cell perimeter

The cell perimeter (*P*) was estimated from the boundary of the segmented mask using the standard perimeter measure implemented in *regionprops*^59^, which approximates the contour length by summing the lengths of pixel edges along the boundary:

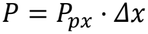

where *P*_px_ is the perimeter in pixels and Δx is the pixel length.

#### Cell circularity

Cell circularity (*C*) was computed as the normalized ratio of the cell projected area (A) to cell perimeter (P), following the standard definition used in image cytometry^61^:

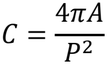

*C* = 1 for a perfect circle and decreases towards zero as the cell shape becomes more elongated or irregular.

#### Cell elongation

Cell elongation (*El*) is defined as the ratio of the major axis length to the minor axis length of the equivalent ellipse fitted to the segmented cell mask, as implemented in *regionprops*^59^:

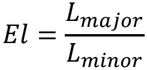

where *L*_major_ and *L*_minor_ are the lengths of the major and minor axes of the region (µm). *El* = 1 indicates a circular cell; values greater than 1 indicate progressive elongation.

#### Cell volume

Because AFM measures only the upper surface of adherent cells, cell volume *V* (µm^3^) was estimated by integrating the height map over the projected cell area — equivalent to the volume enclosed between the upper cell surface and the substrate plane:

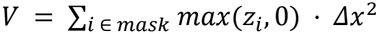

Rare aberrant negative height values were set to zero before summation to avoid artefactual volume subtraction.

Morphometric parameters were exported as a single CSV file containing one row per cell, with all metrics and the segmentation threshold offset used for each cell. Three-dimensional surface plots were rendered using an isometric projection with directional shading computed from the local height gradient. Morphometric parameters and three-dimensional surface plots are available as additional files (see Supplementary Information).

All analyses were performed with a custom open-source Python application built on Flask, NumPy, SciPy and scikit-image^59,62,63^. The application provides a web-based interface for batch analysis, interactive segmentation, manual cell separation, session management, export of results (CSV) and figures (PNG, ZIP).

### Graphs and statistical analysis

All graphs in this work were generated using Origin 2022 (https://www.originlab.com). Statistical analysis and data comparisons were performed using two-tailed Welch’s t-test and one-way ANOVA implemented in GraphPad Prism 11 (https://www.graphpad.com). In both cases, unpaired tests were performed assuming sampling from a normal distribution. In the ANOVA test, multiple comparisons were performed to compare the mean of each column with the mean of any other column using Tukey’s test (^*^, *p*<0.05; ^**^, *p*<0.01; ^***^, *p*<0.001; ^****^, p<0.0001). Residual, homoscedasticity and QQ plots were generated to confirm the main test assumptions and normal distribution of the errors.

## Supplementary Information

Supplementary Information is available from the Wiley Online Library or from the author. It includes the validation of the THP-1 and BMDM differentiation and polarization protocols (Figures S1 and S2), additional morphometric and mechanical data (Figures S3–S6 and S8), the comparison of analytical models for force-curve fitting (Figure S7) and additional comparisons between the THP-1 and BMDM models (Figure S9).

## Acknowledgements

Experiments conducted in this work were funded by the Spanish Ministry of Science and Innovation (BIOMEDSPM4.0, PID2022-142297NB-I00) and the Horizon Europe program under the MSCA doctoral network (SPM4.0, grant agreement No. 101168976). BC acknowledges the Spanish Ministry of Science and Innovation for a predoctoral grant (FPI, PREP2022-000801) and the Generalitat de Catalunya for support through the CERCA programme. GG acknowledges the ICREA foundation (ICREA Academia award). AC acknowledges the Bartolo Longo foundation for the research on cancer (https://fondazionebartololongo.com/), the Institute of Nanoscience and Nanotechnology of the University of Barcelona IN2UB (Ajuts a la Recerca Transversal ART2020) and funding from Academist/Sysmex Diagnostics SE under the project *A touch-based microscope to detect cancer* (https://academist-cf.com/projects/251?lang=en).

## Ethics Approval Statement

Mouse experiments were approved by the CEEA-PCB (Barcelona), under protocol number 24-014 and conducted in accordance with national animal welfare regulations.

## Author Contributions

A.C. and G.G. conceptualized the project’s idea

A.C. and G.G. supervised the project

B.C., L.A., C.C. performed experiments

B.C., A.C., G.G., M.C., S.S. performed data analysis

A.C., B.C., C.C., L.A. prepared the figures

A.C., B.C., L.R., G.G. wrote the manuscript

A.C., G.G., G.B., L.R. acquired funds for the project All authors contributed to research, editing, and approval of the manuscript.

## Data Availability Statement

All data generated or analyses performed in this study are included in this document and the Supplementary information files.

