## Supplementary Information for "Force Volume Atomic Force Microscopy of THP-1 and Primary Mouse Macrophages Reveals that the Mechanical Signature of Macrophage Polarization is Model-Dependent"

### 1. Description of the Repository folder

*Morphometric analysis.zip*

This folder contains the height maps, segmentation masks and 3D views generated for the morphometric analysis of the cells with the typical shape of each phenotype shown in the paper. All morphological parameters obtained are summarized in the CSV file included in each of the two folders (BMDM Mouse and THP-1).

*Raw data and Data analysis 1.zip*

Here, the results of the batch processing routine performed with the spherical model with bottom effect correction are presented, using the geometric contact point for height determination, as described in the Materials and Methods section. Within this folder, data are organized according to the following hierarchical tree structure:

1) Cell model (BMDM Mouse/THP-1/Mouse_M1_whole_cell_analysis)
2) Individual cells are indicated as Model_Experiment_Phenotype_Replicate(if any)_Cell number_Shape (optional). When shape is not indicated in the filename, data refer to the typical shape for that phenotype.
3) Each cell folder contains the raw QI map (.jpk-qi-data) of the specific cell and the results of the analysis, i.e.:
   a) all processed curves with fitting results (‘curves’ folder)
   b) the ROI and substrate cutouts, i.e. indexed pixels belonging to these two regions (obtained from the QI maps), plus the image dimension in pixels (‘cutouts’ folder)
   c) *E*, $R^2$ and Zc vs x, y (‘maps’ folder)
   d) all plots generated from the analysis, like those in Figure 2 and Figure S7 of the manuscript (‘plots’ folder)
   e) the different topographies obtained during the analysis (‘topographies’ folder).
   f) a file with calibration and fitting parameters (log_file.txt)
   g) a summary containing *E*, Zc, $R^2$ and true topography values for the cellular ROI (results_summary.txt).

*All cell maps - Panel.zip*
Each file in this folder is a single image panel showing the optical images, true topography, *E* and $R^2$ maps of a group of cells of the same model, phenotype and shape, as indicated in the filename (larger groups are split into several files).

*Figures data.zip*
The "Figures data" folder is organized into one subfolder per figure and contains the source data for the following panels in .csv format:

- Main figures: Figure 1e–l; Figure 2a, c–e, g–j; Figure 3b–j; Figure 4b–j; Figure 5d–i; Figure 6d–i; Figure 7a–c.
- Supplementary figures: Figure S1b–d; Figure S2b–e; Figures S3–S6; Figure S7a–e, g–j; Figure S8b–d, f–h; Figure S9.

The Repository folder is available upon request at: https://doi.org/10.5281/zenodo.23888707.

### 2. Validation of sample preparation protocols of THP-1 and BMDM macrophages

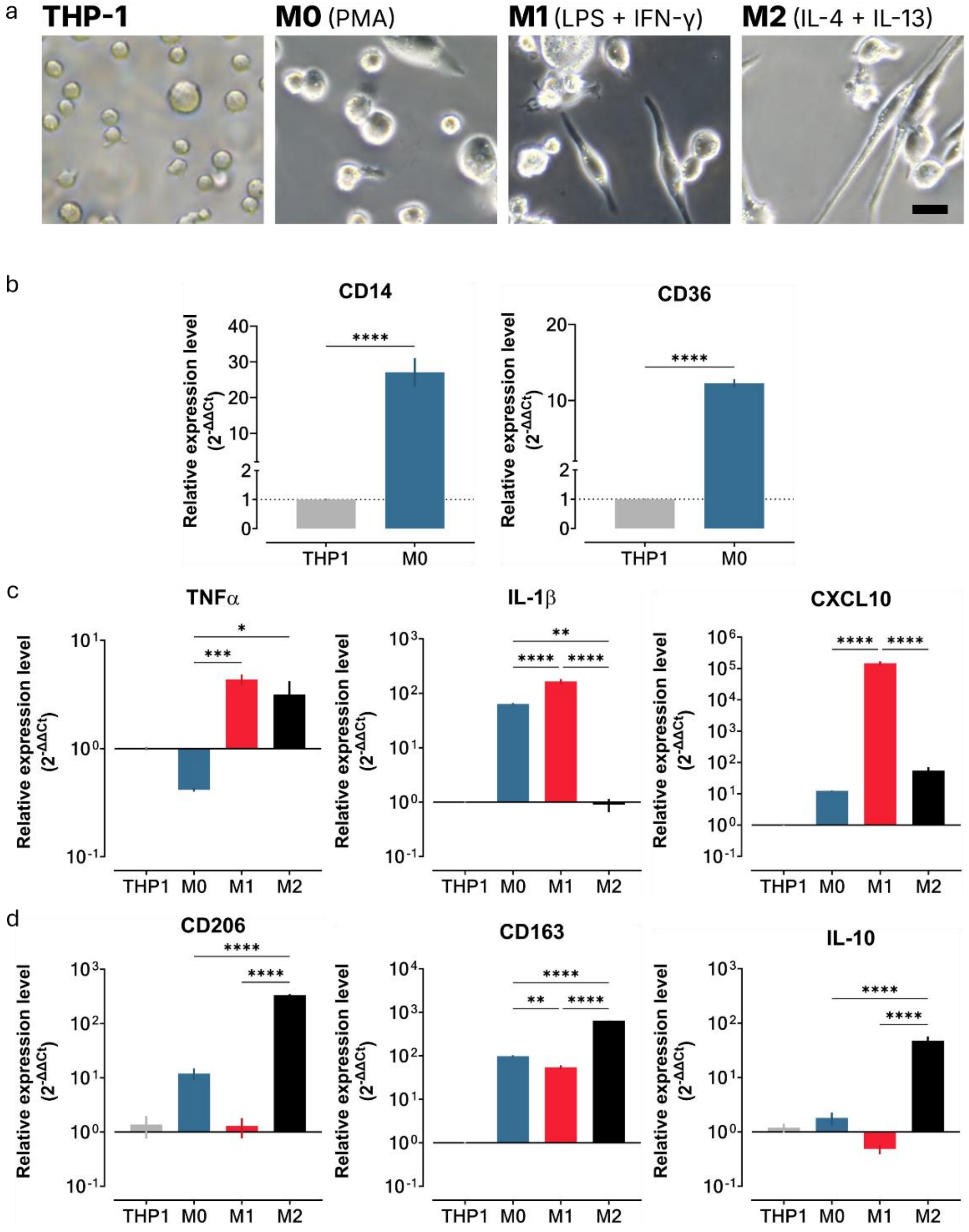


**Figure S1. Differentiation and polarization of THP-1 macrophages.** (a) Representative bright-field images of human THP-1 monocytes and THP-1-derived macrophages (scale bar: 20 µm, same for all images). PMA stimulation (see the article's Materials and Methods) induces the differentiation of suspended THP-1 monocytes into adherent macrophages (M0). M0 cells are then polarized into pro-inflammatory (M1; LPS+IFN-γ) or anti-inflammatory (M2; IL-4+IL-13) macrophages, see the article's Materials and Methods. (b) mRNA expression of monocyte-to-macrophage differentiation markers: the monocyte marker CD14 (left) and the macrophage marker CD36 (right), ****p<0.0001. (c) mRNA expression of pro-inflammatory macrophage markers: TNF-α (left), IL-1β (middle) and CXCL10 (right) (*p<0.05, **p<0.01, ***p<0.001, ****p<0.0001). (d) mRNA expression of anti-inflammatory macrophage markers: CD206 (MRC1) (left), CD163 (middle) and IL-10 (right) (**p=0.0025, ****p<0.0001). In (b–d), mRNA levels were measured by RT-qPCR and quantified with the 2^(−ΔΔCt) method, normalized to GAPDH and expressed relative to undifferentiated THP-1 monocytes [1]. Data are shown as mean ± SEM (n = 6 replicates from two independent experiments).

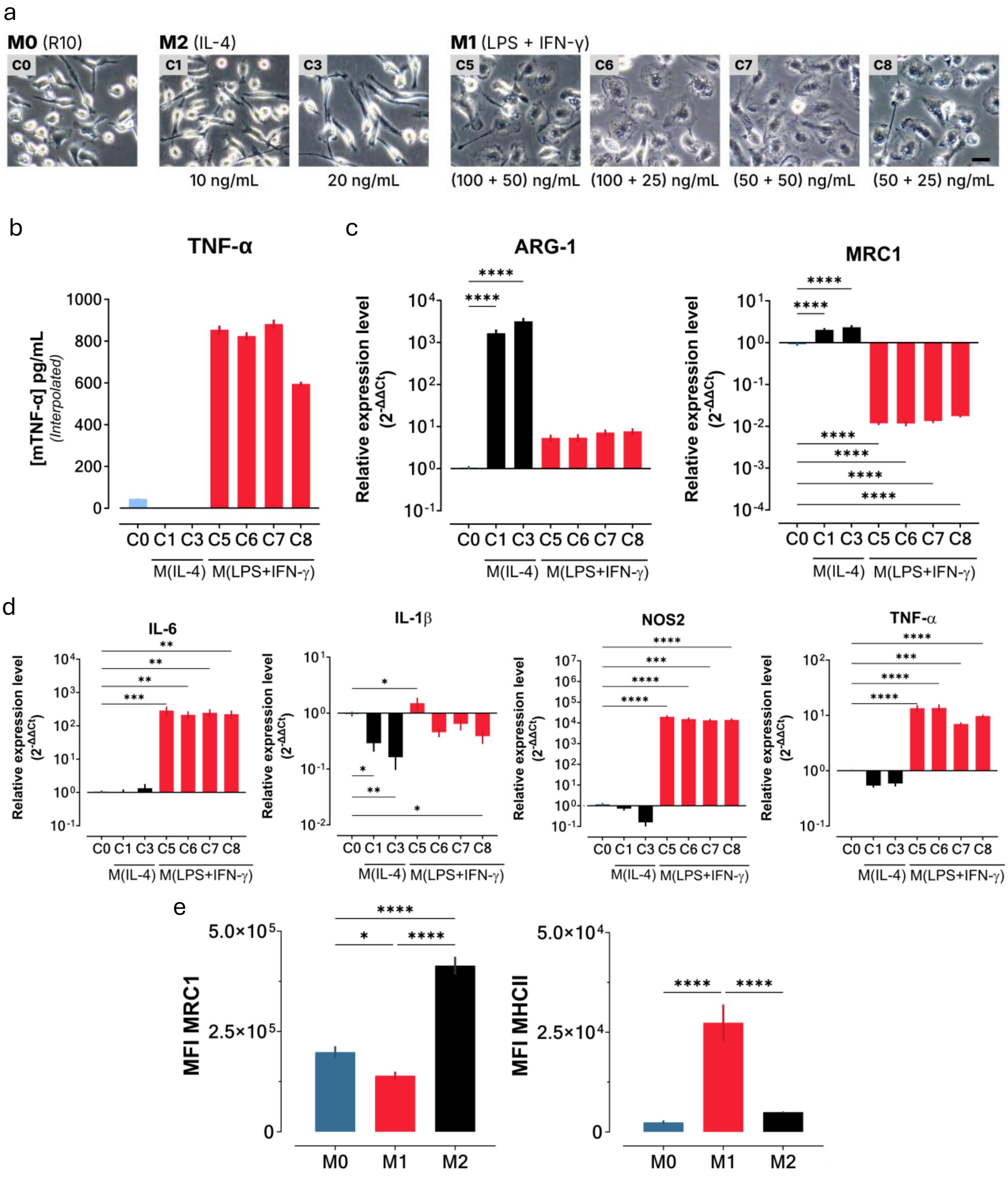


**Figure S2. Differentiation and polarization of mouse bone marrow-derived macrophages (BMDMs).** (a) Representative bright-field images of untreated BMDMs (C0, M0) and BMDMs stimulated with different concentrations of IL-4 or LPS + IFN-γ to induce their polarization into anti-inflammatory (M2) or pro-inflammatory (M1) macrophages, respectively. Conditions: C0, control (R10 medium only); C1, IL-4 (10 ng/mL); C3, IL-4 (20 ng/mL); C5, LPS + IFN-γ (100 + 50 ng/mL); C6, LPS + IFN-γ (100 + 25 ng/mL); C7, LPS + IFN-γ (50 + 50 ng/mL); C8, LPS + IFN-γ (50 + 25 ng/mL). Scale bar: 20 µm, same for all images. (b) TNF-α concentration (pg/mL) in BMDM culture supernatants after 24 h of stimulation under each condition (C0, C1, C3, C5–C8), quantified by ELISA. Bars represent mean ± SEM (N = 6; 3 female and 3 male). (c) mRNA expression of the anti-inflammatory markers ARG-1 (left) and CD206 (MRC1) (right) after 24 h of stimulation (****$p < 0.0001$). (d) mRNA expression of the pro-inflammatory markers IL-6, IL-1β, NOS2 and TNF-α (left to right) after 24 h of stimulation (*p<0.05, **p<0.01, ***p<0.001, ****p<0.0001). mRNA levels in (c, d) were measured by RT-qPCR and quantified with the 2^(−ΔΔCt) method [1], normalized to β-actin and expressed relative to the control (C0). Bars represent mean ± SEM (N = 6; 3 female and 3 male). (e) Median fluorescence intensity (MFI) of the surface markers MRC1 (CD206) and MHCII, measured by flow cytometry in M0, M1 and M2 BMDMs, using the stimulation conditions reported in this article (C3 and C5). M2 macrophages showed robust MRC1 (CD206) expression, confirming an immunoregulatory phenotype. M1 macrophages showed significantly higher MHCII expression than M0 and M2 cells, consistent with enhanced antigen presentation in the pro-inflammatory state. Data represent mean ± SEM (N = 6; 3 female and 3 male; n = 3 per animal) (*p<0.05, ****p<0.0001).

### 3. Additional data

#### 3.1 Additional data for Figure 1

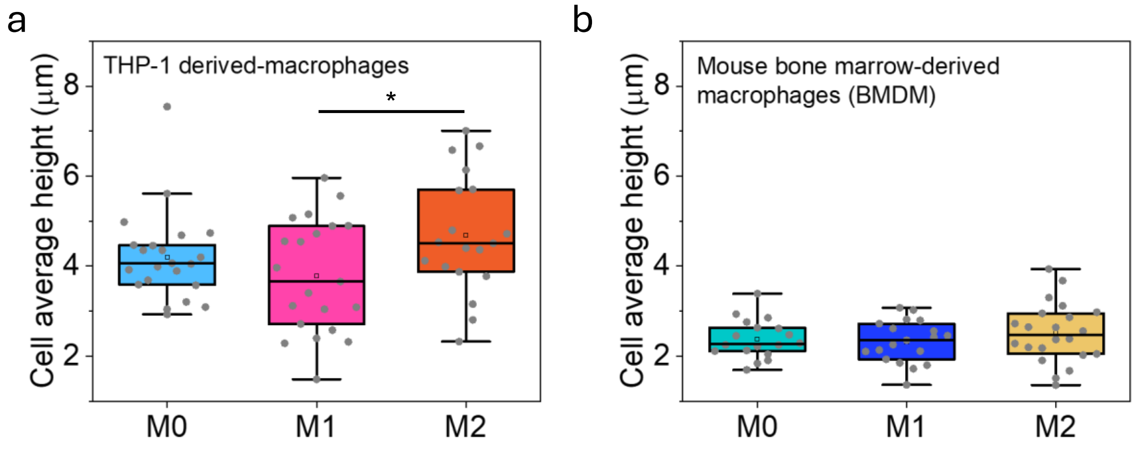


**Figure S3.** Additional results from the morphometric analysis. Average cell height extracted from the cells' segmentation for (a) THP-1-derived macrophages and (b) BMDM. The ROIs used for this analysis are the same as those shown in Figure 1 of the main text and are collected in the repository folder "Morphometric analysis".

#### 3.2 Additional data for Figures 3 and 4

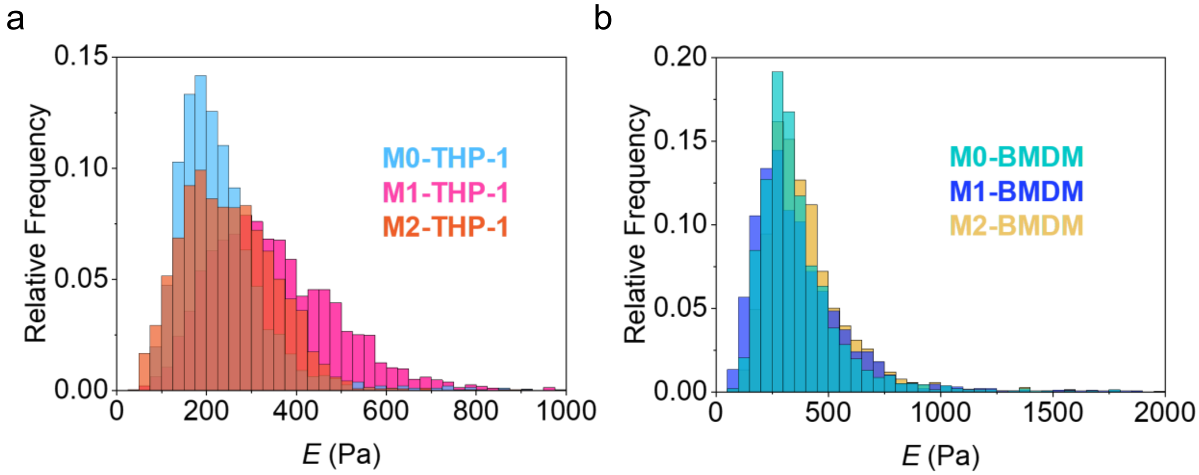


**Figure S4.** Distribution histograms of *E* data, obtained using all image pixels belonging to cells with the typical shapes associated with M0, M1 and M2 phenotypes (see Figures 3 and 4 of the main text). Data are separately represented for the THP-1 (a) and the BMDM model (b). In both cases, the *E* distribution is log-normal.

### 3.3 Additional data for Figure 5

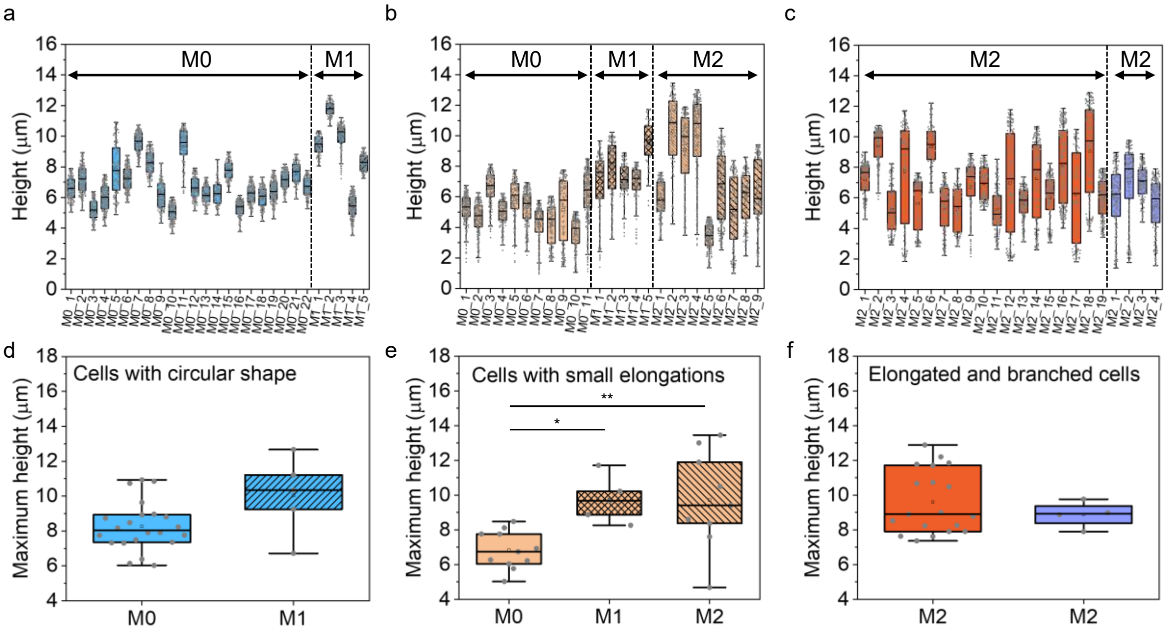


**Figure S5.** (a, b) Height of single macrophages belonging to the THP-1 model with a circular shape (these cells were mainly present in M0 and M1 samples) (a) and small elongations (these cells were present in all samples) (b). (c) Height of elongated cells and branched cells with dendritic shape present in M2 samples. (d–f) Corresponding maximum height per cell, with the results of statistical analysis, i.e. Welch's t-test (d and f) and ANOVA (e).

### 3.4 Additional data for Figure 6

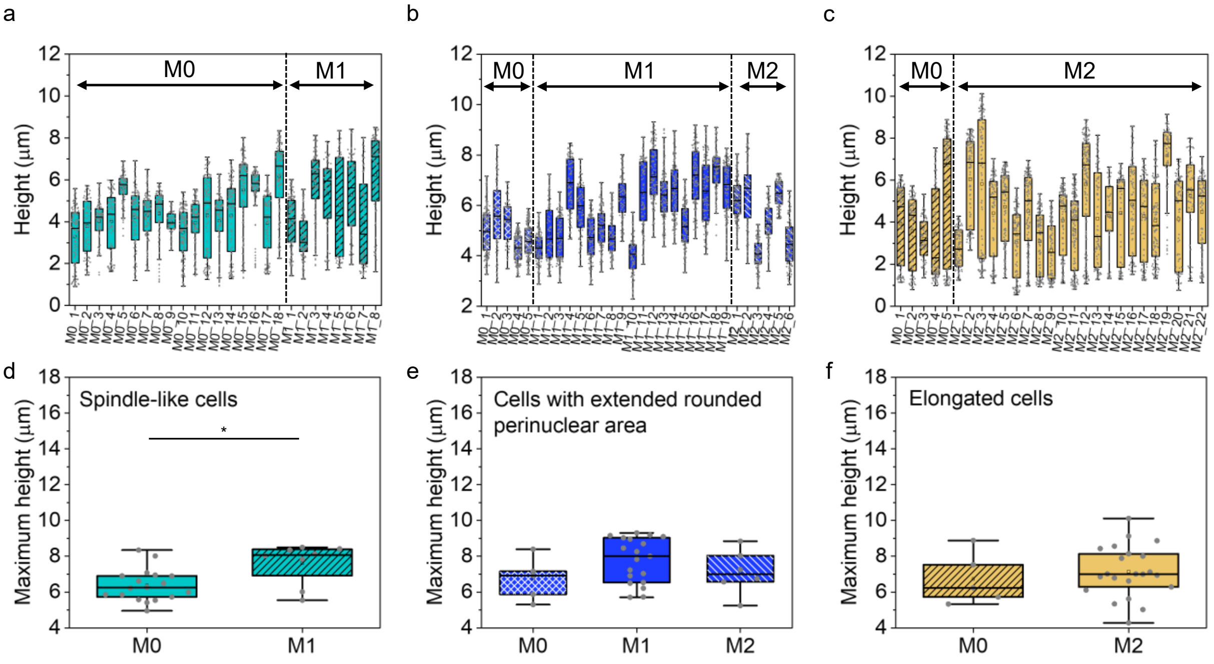


**Figure S6.** (a–c) Height of single macrophages belonging to the BMDM model: spindle-like cells (these cells were mainly present in M0 and M1 samples) (a), circular shape with extended rounded perinuclear area (these cells were present in all samples) (b), and very pronounced elongations (longer than 50 µm), present in M0 and M2 samples (c). (d–f) Corresponding maximum height per cell, with the results of statistical analysis, i.e. Welch's t-test (d and f) and ANOVA (e).

#### 3.5 Comparison among mechanical models

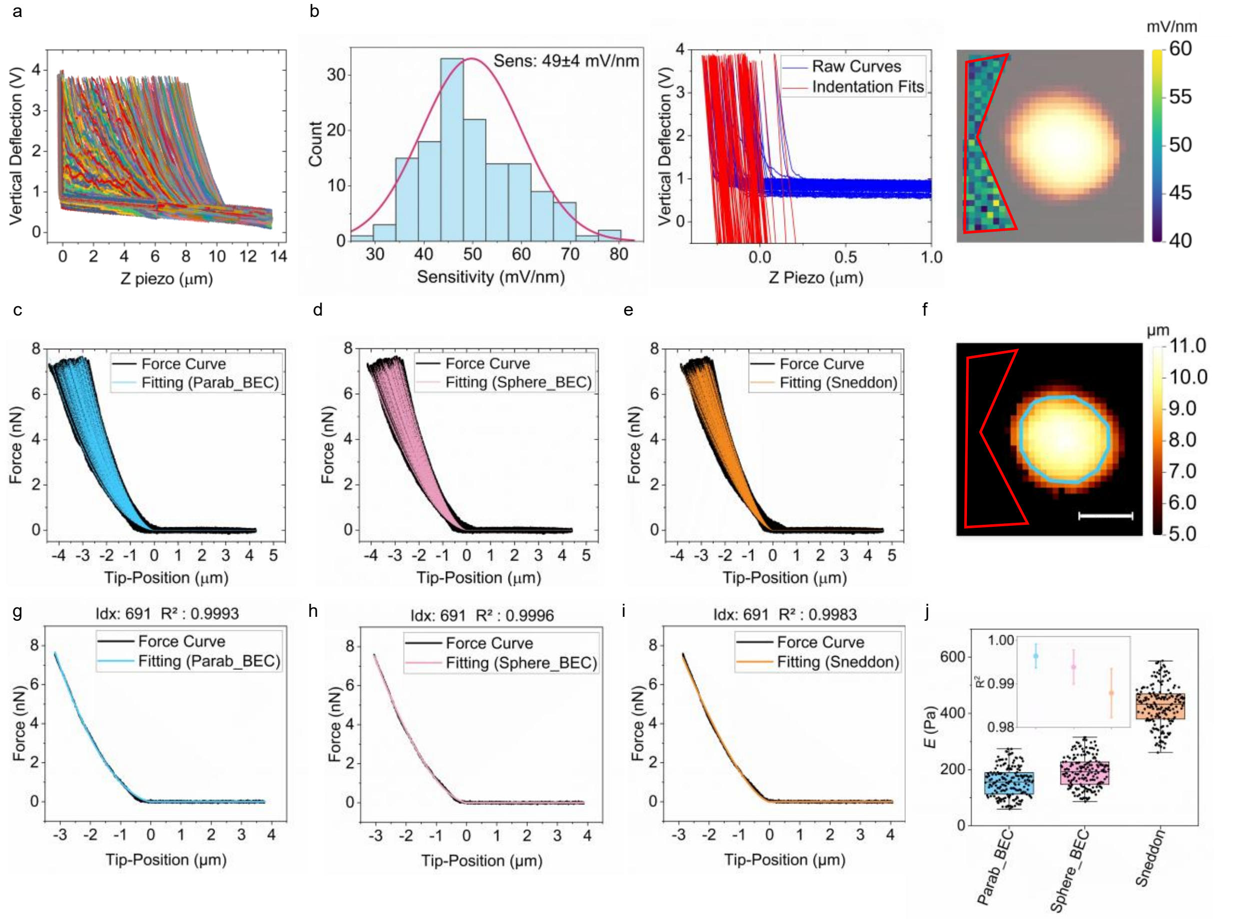


**Figure S7.** Mechanical analysis of an M0 macrophage from THP-1 performed with different analytical models. (a) Raw curves; (b) sensitivity analysis using the substrate region highlighted in the topographic image; (c–e) baseline-corrected, calibrated force vs. tip-position curves (in black) with fitted curves overlaid. Fits are performed using the model of a paraboloid indenter with bottom effect correction (BEC) (curves in light blue in (c)) [2], a spherical indenter with BEC (curves in pink in (d)) [2], [3] and a spherical indenter without BEC (Sneddon's model, curves in orange in (e)). Experimental curves are shifted to the fitted contact point (see also Materials and Methods). (f) True topography image (scale bar = 10 µm). (g–i) Individual curve fitted with the three models with curve index and fit $R^2$ indicated. (j) $E$ values for the cellular ROI obtained with the three models. (j, inset) Average $R^2$ from the fits in the cellular ROI ($R^2$=0.996 ± 0.003, 0.994 ± 0.004 and 0.988 ± 0.006 for the fit with the paraboloid model with BEC, the spherical model with BEC and the Sneddon model without BEC, respectively). All data and plots obtained with the Mechanotool custom software are provided as additional files (see Section 1, Description of the Repository folder).

#### 3.6 Mechanical analysis of M1 macrophages from BMDM including large, roughly circular perinuclear areas

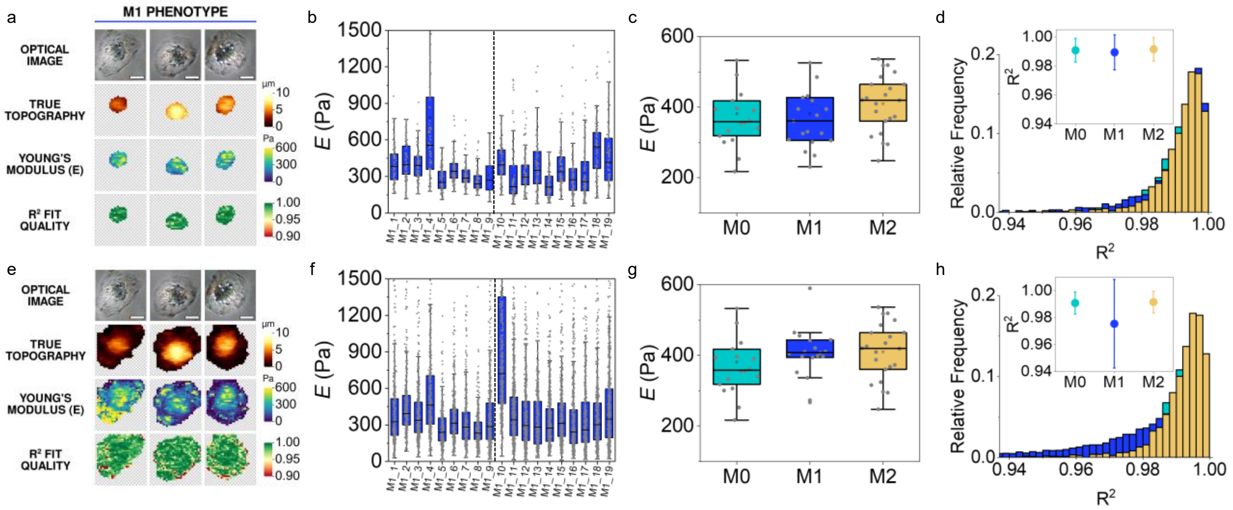


**Figure S8.** Comparison of the Young's modulus results obtained in primary M1 macrophages (BMDM) when the large perinuclear cell areas are included in the analysis. (a–d) Analysis restricted to the nuclear region, as in the main text (Figure 4). (e–h) Analysis including the whole cell area, i.e. the large perinuclear areas. (a, e) Optical images, true topography, E and $R^2$ maps of three selected cells (the same cells are shown in (a) and (e); scale bar = 10 µm). (b, f) E values of all individual M1 cells analyzed (the vertical line separates the experimental days). (c, g) Average E per cell for the M0, M1 and M2 phenotypes. (d, h) Normalized distribution histograms of $R^2$ values including all image pixels, with average values for each phenotype (insets). When the whole cell area was included, the fitting accuracy of M1 cells was poorer ($R^2 = 0.975 \pm 0.033$) and the Young's modulus tended to be slightly higher, but the comparison test (ANOVA) showed no significant differences among the three phenotypes.

#### 3.7 Additional data for Figure 7

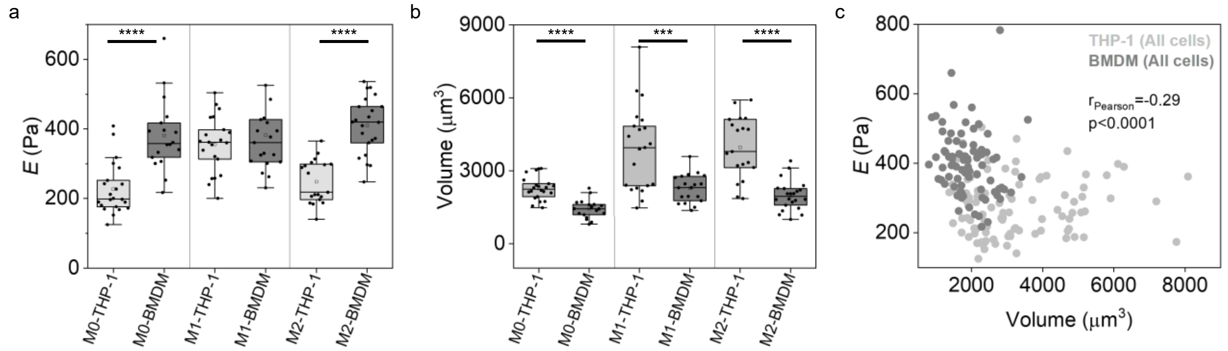


**Figure S9.** (a) Young's modulus values highlighting the differences between the THP-1 and BMDM cell models. Vertical lines separate the M0, M1 and M2 phenotypes. (b) Corresponding plot of the cellular volume from morphometric analysis. In both graphs the results of comparison with Welch's t-test are included for statistically different samples. (c) *E* vs volume with the results of the correlative analysis considering all cells from THP-1 and BMDM as a single group and all cellular shapes.
